# Genotypic and Phenotypic Analyses Show *Ralstonia solanacearum* Cool Virulence is a Quantitative Trait Not Restricted to “Race 3 biovar 2”

**DOI:** 10.1101/2024.06.13.598915

**Authors:** Ronnie J. Dewberry, Parul Sharma, Jessica L. Prom, Noah A. Kinscherf, Tiffany Lowe-Power, Reza Mazloom, Xuemei Zhang, Haijie Liu, Mohammad Arif, Michael Stulberg, Lenwood S. Heath, Kellye Eversole, Gwyn A. Beattie, Boris A. Vinatzer, Caitilyn Allen

## Abstract

Most *Ralstonia solanacearum* species complex strains cause bacterial wilts in tropical or subtropical zones, but the group known as Race 3 biovar 2 (R3bv2) is cool virulent and causes potato brown rot at lower temperatures. R3bv2 has invaded potato-growing regions around the world but is not established in the United States. Phylogenetically, R3bv2 corresponds to a subset of the *R. solanacearum* phylotype IIB clade, but little is known about the distribution of the cool virulence phenotype within phylotype IIB. Therefore, genomes of 76 potentially cool virulent phylotype IIB strains and 30 public genomes were phylogenetically analyzed. A single clonal lineage within the sequevar 1 subclade of phylotype IIB that originated in South America has caused nearly all brown rot outbreaks worldwide. To correlate genotypes with relevant phenotypes, we quantified virulence of ten *Ralstonia* strains on tomato and potato at both 22°C and 28°C. Cool virulence on tomato did not predict cool virulence on potato. We found that cool virulence is a quantitative trait. Strains in the sequevar 1 pandemic clonal lineage caused the most disease, while other R3bv2 strains were only moderately cool virulent. However, some non-R3bv2 strains were highly cool virulent and aggressively colonized potato tubers. Thus, cool virulence is not consistently correlated with strains historically classified as R3bv2 group. To aid detection of sequevar 1 strains, this group was genomically delimited in the LINbase web server and a sequevar 1 diagnostic primer pair was developed and validated. We discuss implications of these results for the R3bv2 definition.

## Introduction

The *Ralstonia solanacearum* species complex (RSSC) is a monophyletic but heterogenous group of plant pathogenic bacteria that cause lethal wilt diseases of many crops by colonizing the water-transporting xylem vessels (Hayward 1991). The thousands of genetically distinct strains in the RSSC infect a broad range of plants and are found on six of the seven continents (Denny 2006). Comparative genomic analyses have divided the RSSC into four phylotypes (phyl.) that have distinct geographic origins: phyl. I (Asia) and III (Africa) correspond to the named species *R. pseudosolanacearum*; phyl. II (the Americas) corresponds to the named species *R. solanacearum* and is sub-divided into sub-phyl. IIA, IIB, and IIC; and phyl. IV (Indonesia and Japan), also known as *R. syzygii* (Safni et al. 2014; P. Sharma et al. 2022; Fegan M. 2005; Yabuuchi et al. 1995) (**Figure 1A**). Within phylotypes, RSSC strains are further sub-divided into sequevars (seq.) based on the partial sequence of the endoglucanase (*egl*) gene (Fegan M. 2005). We recently used a reverse ecology approach to assign strains in the RSSC to population clusters based on recent recombination events; this whole-genome analysis method largely recapitulated the sequevar subclassification and confirmed its general utility (P. Sharma et al. 2022).

**Figure 1.**
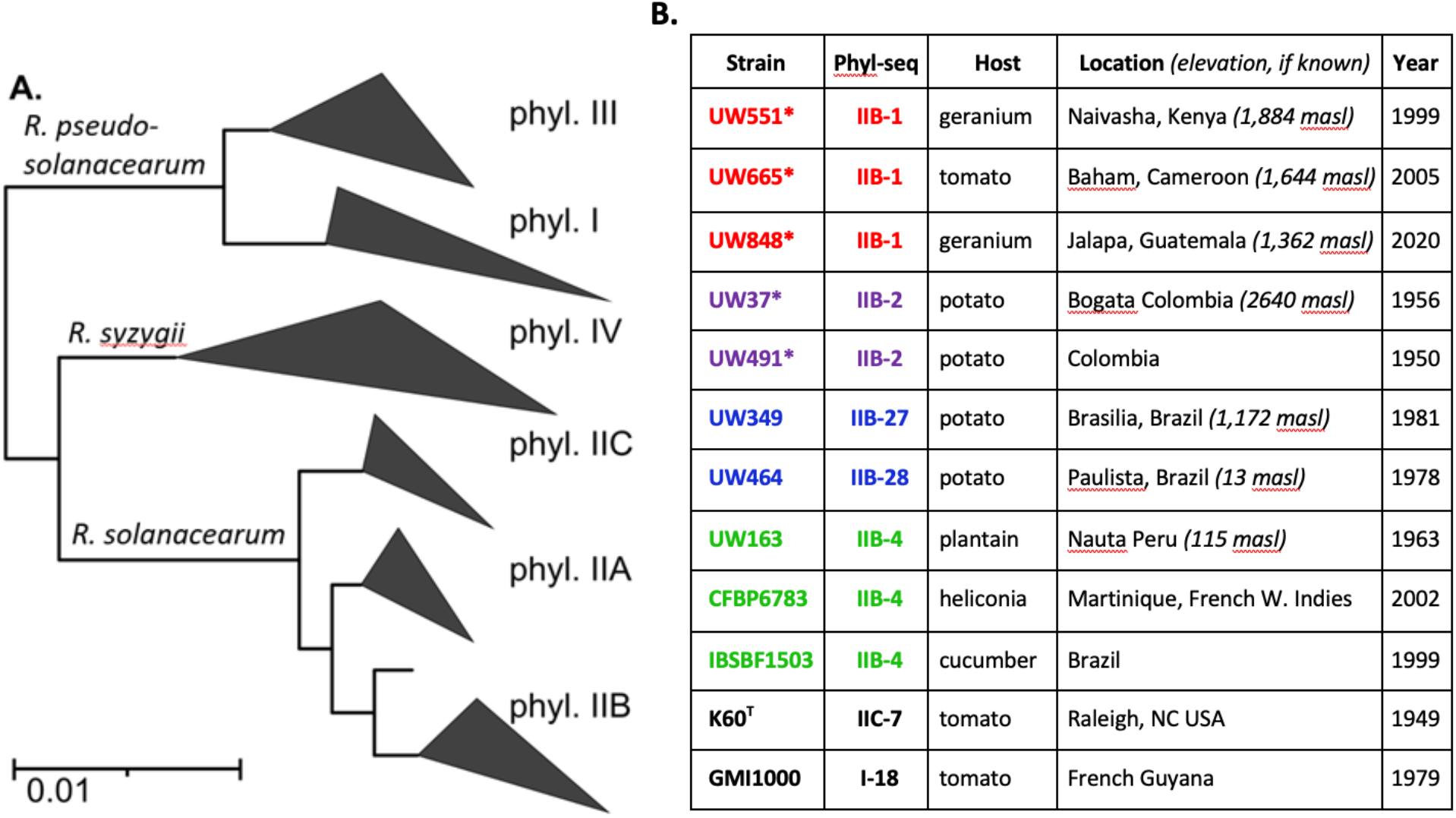
*Ralstonia solanacearum* species complex (RSSC) phylogenetic tree and strains used in phenotyping assays. A. Phylogenetic tree showing relationships among the phylotypes of the RSSC, based on 49 conserved bacterial genes using 466 public genomes. B. Characteristics of *Ralstonia* strains used in phenotyping assays. Phyl-seq: each strain’s phylotype and sequevar classification were respectively determined by the phylotype multiplex PCR and *in silico* PCR and comparative analysis of an internal fragment of the conserved *egl* gene (see Methods). Phylotype IIB strains are color-coded here and in other figures as follows: red, sequevar (seq.). 1; purple, seq. 2; blue, seq. 27 and 28; green, seq. 4; black, non-phylotype IIB strains. Asterisks indicate strains belonging to seq. 1 and 2, which are considered R3bv2; superscript T indicates that K60 is the *R. solanacearum* type strain.

Bacterial wilt is primarily a problem for growers in tropical to warm-temperate zones, but some strains in phyl. IIB cause disease in tropical highlands where temperatures are lower (Champoiseau, Jones, and Allen 2009; Allen, C., Kelman, A., French, E. R 2001). This phenotype of inducing symptoms at low temperatures (at or below 22°C) is known as cool virulence. Cool virulent phyl. IIB strains cause potato brown rot, a destructive disease that seriously threatens food security in parts of South and Central America, Africa, and Asia (Elphinstone 2005). They also occasionally cause disease in temperate zones of Europe and Asia (Parkinson et al. 2013). The highest genetic diversity of potato brown rot strains is found in the Andean highlands, which is also the center of diversity for potato. Andean potato brown rot strains are a moderately diverse group that belong to seq. 1, 2, 25, 27, and 28 (Fegan M. 2005). Most of these strains are not found beyond South America. Epidemiological and whole-genome data indicate that a single clonal lineage of seq. 1 has spread around the world out of South America (Clarke et al. 2015). We call this the Brown Rot Pandemic Lineage (hereafter BRPL).

Since *R. solanacearum* strains were historically described based on supposed host range (race) and metabolic traits (biovar) (Denny 2006), potato brown rot strains are widely referred to as *R. solanacearum* Race 3 biovar 2 (R3bv2) in the literature and in biosecurity regulations. Several molecular tests detect seq. 1 and 2 (Stulberg and Huang 2015; Stulberg et al. 2018) or the larger group of Andean brown rot sequevars (Weller et al. 2000; Fegan et al. 1998). However, a relatively small number of *R. solanacearum* genomes were available when these molecular tests were developed. Additional evaluation using a larger number of genomes would be beneficial to verify accuracy of test results.

R3bv2 has been repeatedly introduced into the United States on ornamental geranium cuttings (Kim et al. 2003; Roman-Reyna et al. 2021; D. L. Strider, R. K. Jones, R. A. Haygood 1981; Williamson et al. 2002), most recently in the spring of 2020 (Roman-Reyna et al. 2021). Despite these multiple introductions, R3bv2 is not established in the United States. Although the primary threat posed by R3bv2 is believed to be their cool virulence, that is, their potential to cause disease in the northern potato-growing regions of the country, it is not known if all strains that belong to seq. 1 and 2 are cool virulent or if cool virulence is limited to these sequevars. Pioneering studies by Bocsanczy, Norman, and co-workers (Bocsanczy et al. 2012;)Norman et al. 2009; Bocsanczy, Huguet-Tapia, and Norman 2017) suggested otherwise. They found that some *R. solanacearum* phyl. IIB strains belonging to seq. 4, established in the state of Florida, can infect and wilt tomato and potato plants at the very low temperature of 18°C.

The objective of this work was to further test the strength of the correlation between cool virulence and membership in *R. solanacearum* seq. 1 and 2 to inform regulation and risk assessment. To this end, we first used comparative evolutionary genomics to define the precise phylogenetic relationships among strains in seq. 1 and 2 and other potentially cool virulent phyl. IIB strains from South America. We then measured the virulence of representative strains on tomato and potato plants at 22°C and 28°C. Lastly, by correlating the resulting phenotypic and phylogenomic data, we attempted to circumscribe the authentically threatening strains based on genome sequence data. We found that while strains from the globally distributed BRPL of seq. 1 generally had the highest virulence at 22°C on potato, cool virulence is a quantitative trait that varies among strains inside and outside of the currently regulated R3bv2 group. Further, the bacterial response to temperature interacts with host preference, meaning that cool virulence on tomato did not predict a strain’s ability to wilt potato at 22°C. We discuss the implications of our results for regulation of *R. solanacearum* and provide the relevant genomic and phenotypic data in LINbase (Tian et al. 2020) to facilitate genome-based identification of putative cool virulent strains.

## Methods

### Bacterial strains, DNA extraction, genome sequencing, and genome assembly

Thirty *Ralstonia* strains with genome sequences publicly available in 2021 and 76 strains selected for genome sequencing are listed in **Supplementary Table 1** and summarized in **Table 1**, along with their metadata, including phylotype, sequevar, host of isolation, country of isolation, state, region or province of isolation (when available), and date of isolation (when available). To obtain DNA for genome sequencing, strains were cultured in rich casamino acid-peptone-glucose (CPG) medium as previously described with minor adjustments (Khokhani et al. 2018). Briefly, a single colony from a fresh CPG agar plate was used to inoculate a 5-ml overnight culture in CPG broth. Genomic DNA was extracted from 1 ml of overnight culture using the Epicentre MasterPure DNA extraction kit according to the manufacturer’s instructions. The initial lysis step of this protocol is a validated inactivation protocol for regulated R3bv2 *R. solanacearum* strains (Hayes et al 2022). Sequencing on the PacBio Sequel II platform was performed at the Genomics Resource Center at the University of Maryland using a 48-plex multiplexed library with size selection and a single 8M SMRT cell. Sequencing on the Illumina platform was performed at the DNA Facility at Iowa State University using one Next-Tera XT 96-multiplexed library kit (Illumina) and a single Illumina HiSeq 3000 lane. PacBio genomes were assembled using Canu (version 2.0) (Koren et al. 2017); Illumina genomes were assembled using SPAdes (version 3.14.0) (Prjibelski et al. 2020). All genome assemblies were assessed for quality using the CheckM (version 1.0.13) tool (Parks et al. 2015).

**Table 1.**
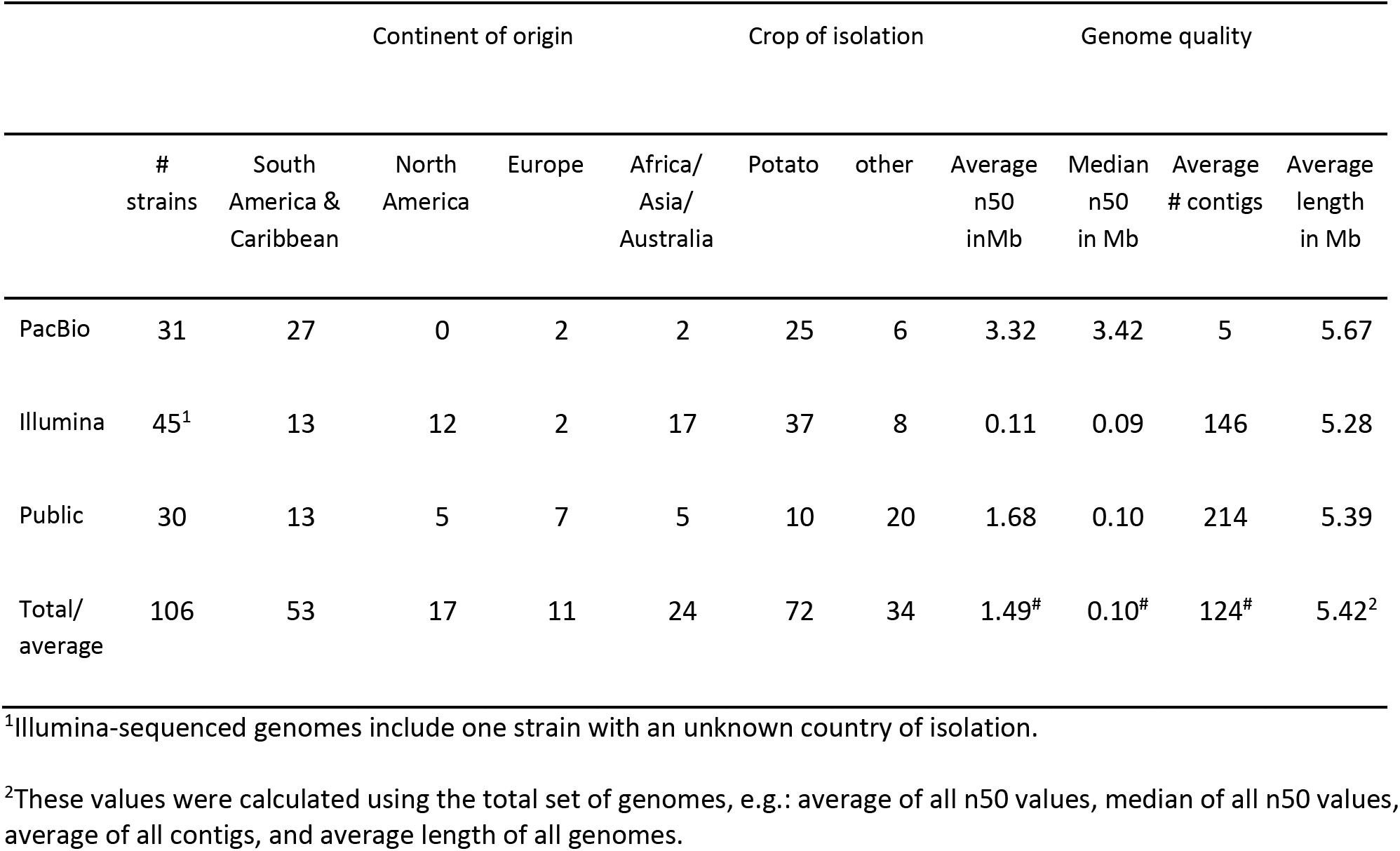
Summary of strain metadata and genome quality.

### Pangenome analysis and construction of core genome phylogeny

A pangenome analysis of the genomes was performed using PIRATE (version 1.0.4) (Bayliss et al. 2019). Genomes were annotated using BAKTA (version 1.5.0) (Schwengers et al. 2021) with default settings. Annotated genomes were used as input for PIRATE, which provided a core-gene alignment by using the following parameters: –a to get a multiFASTA core-gene alignment file as output and –k for faster homology searching with the ––diamond option specified. Gene families in the core-gene alignment file were then used to create a core-gene alignment file for input into IQtree (version 2.0.3) (Minh et al. 2020) using automated model-selection to obtain a maximum-likelihood (ML) phylogenetic tree. The final phylogenetic tree was visualized using the ggtree (Yu et al. 2017) package in R. The PIRATE output file, with all gene families, was further used to find differences in gene content between phylogenetic clades. For clades with higher representation (more than or equal to 20 strains), a gene was considered core to the clade if more than 95% of the strains in that clade had that gene. For clades with lower representation (less than 20 strains), genes present in all but one strain were considered core to the clade. A presence and absence matrix was constructed for all clades with a score of 1 if a gene was considered to be core to that clade and a score of 0 otherwise. The resulting matrix was used as input to construct an Upset plot using the UpSetR (Conway, Lex, and Gehlenborg 2017) package in R.

### Construction of a 49-gene tree for publicly available RSSC genomes

As described (Lowe-Power et al. 2022), we uploaded all publicly available RSSC genomes (n=466), including the genomes from this study, to a public KBase narrative (Arkin et al. 2018). The KBase SpeciesTree tool was used to build a FastTree-based phylogenetic tree from 49 conserved genes (Price, Dehal, and Arkin 2010). The newick tree file was uploaded to iTol (Letunic and Bork 2021), and the major clades were collapsed.

### Single Nucleotide Polymorphism (SNP) detection and tree construction

SNP analysis was performed using the automated SNP detection CFSAN-snp-pipeline tool (version 2.2.0) (Davis et al. 2015) with default settings. Reads of all genomes were aligned against a chosen high-quality reference genome, CIP10_UW477. For published genomes for which reads were not available, individual reads were simulated using the ART-next generation read simulator tool (version 2.5.8) (Huang et al. 2012). The resulting multi-fasta alignment was then used as input for IQ-TREE (version 2.0.3) (Minh et al. 2020) using automated model-selection to obtain an ML tree. The final SNP tree was visualized using the ggtree (Yu et al. 2017) package in R. The rooted tree was then used as input for the online tool Phylostems (Doizy et al. 2023), which computes temporal signals for molecular tip-dating.

### Average nucleotide identity (ANI) analysis, LIN assignment, and LINgroup circumscription

Pairwise average nucleotide identity (ANI) was measured using pyANI (version 0.2.10) (Pritchard et al. 2015) with default settings. All sequenced strains belonging to phyl. IIB were used for this analysis. The resulting matrix was used to construct a heatmap of ANI values using the function heatmap.2 under the gplots package (Warnes et al. 2015) in R. Public genome assemblies and newly sequenced genome assemblies were uploaded together with strain metadata from **Supplementary Table 1** to LINbase at linbase.org (Tian et al. 2020). Life Identification Numbers (LINs) were automatically assigned and groups of genomes were circumscribed, named, and described as LINgroups.

### Sequevar analysis and virtual PCR

Sequevar assignments were made as described previously (P. Sharma et al. 2022) using a custom bash script that takes a query genome sequence as input and compares it to a database of *egl* gene sequences using BLAST (Camacho et al. 2009). The database was provided by E. Wicker (Wicker et al. n.d.). Sequevars were assigned based on the best hit with 99-100% alignment.

Virtual PCR results for primers 630/631 (Fegan et al. 1998), RSCV (Stulberg and Huang 2015) and RsSA3 (Stulberg et al. 2018) were obtained using a custom script that compares each input genome against the respective PCR product whereby the sequence of the PCR products were obtained by mapping the primers to the genome sequence of UW551 (GCA_002251655.1), a genome known to give a positive PCR result for all three primer sets.

### Cool virulence and potato colonization assays

Inoculum for virulence assays was prepared by adding 1 ml of an overnight *R. solanacearum* culture grown from a single colony in CPG broth as described above to a 500-ml flask containing 300 ml of CPG medium and shaking at 200 rpm at 28°C for 24 h. Bacteria were pelleted by centrifugation, washed by resuspension in water, and adjusted to an optical density at 600 nm (OD_600_) of 0.1, corresponding to ∼1×10^8^ CFU/ml. The exact inoculum density used in each experiment was verified using serial dilution plating.

Tomato virulence assays were conducted as previously described (Khokhani et al. 2018). Briefly, unwounded 21-day-old tomato plants (wilt-susceptible cv. Bonny Best), grown in 4-inch pots at 22°C or 28°C, were inoculated via soil drenching with 50 ml of a 10^8^ CFU/ml bacterial suspension. Disease progress was rated daily using a 0-4 disease index rating scale over 21 days for plants grown at 22°C and over 14 days for those at 28°C. Potato virulence assays were conducted on unwounded 28-day-old potato plants (cv. Russet Norkotah) grown from *in vitro*-propagated cuttings in 8-inch pots at 22°C or 28°C and inoculated using the same soil-soak inoculation protocol as for tomato plants. Potato disease progress was assessed every other day for 40 days at 22°C and 21 days at 28°C, using the same 0-4 rating scale. Virulence assays were independently replicated 3 to 8 times per strain, with each replicate experiment containing 6 plants per strain for both temperatures and for both tomato and potato plants.

Potato colonization ability was assayed on Russet Norkotah potato plants grown at 22°C and inoculated as described for potato virulence assays. Plants were harvested at the first sign of wilt development, which occurred 2 to 3 weeks post inoculation. One stem sample and the two largest tubers were collected from each plant. Potato stems and tubers were surface sterilized for 1 minute in 10% bleach, 5 minutes in 70% ethanol, rinsed with water, and air-dried for 30 minutes. A 0.1 g cross-section of crown tissue or a 0.1 g sample of tuber tissue harvested with a 1.0-mm diameter corerer from the stolon end were added to a 2 ml screw-cap microcentrifuge containing 900 µl of water and ten metal beads. Tissues were homogenized with the Powerlyzer™ 24 (MoBio Laboratories, Inc.) using the following program: Speed=2200, Time=1:30 min, Delay=4 min, Cycle=4. Homogenates were serially dilution plated onto modified SMSA agar plates (“EPPO Diagnostic Protocol for Ralstonia Solanacearum Species Complex” n.d.) without polymyxin B sulfate to determine bacterial cell densities in plant tissues. Colonization assays were independently replicated 3 to 4 times per strain, with each replicate containing data from 6 crowns and 12 tubers per strain.

Disease progress curves were generated for each strain on potato and tomato plants at both 22°C and 28°C. The Area Under the Disease Progress Curve (AUDPC) for each plant and the resulting data for each strain-host combination were subjected to Two-way Repeated Measures Analysis of Variance (Two-way RM-ANOVA). Potato colonization data were analyzed either using One-way ANOVA with Student’s *t* test to test for differences in bacterial population sizes or with Pearson Chi square with odds ratio test to test for frequency of colonization. Statistical significance among strains or treatments was determined based on *P*<0.05 unless otherwise indicated. All phenotypic data were analyzed using the Prism 8 software (GraphPad, San Diego, CA).

## Results

### Genome sequencing and assembly

We assembled the genomes of 31 phyl. IIB strains from long reads obtained on the PacBio platform and genomes of 45 phyl. IIB strains from short paired-end Illumina reads. **Supplementary Table 1** lists the quality of the assemblies for each newly sequenced genome and all the previously available genome sequences that were used in this study. **Table 1** provides a short summary of these results. Raw sequencing data were submitted to the NCBI SRA database and assembled genomes were submitted to the NCBI Assembly database under NCBI Project ID PRJNA775652. Assembled genome sequences and metadata were also uploaded to LINbase and assigned LINs (Life Identification Numbers) reflecting their reciprocal genome similarity (Tian et al. 2020). All genomes shared the LIN prefix 14_A_1_B_0_C_0_D_0_E_3_F_0_G_0_H_ confirming their identity as members of phyl. IIB with a minimum of 97% pairwise ANI (P. Sharma et al. 2022).

### Core genome phylogeny of IIB strains

We began unraveling the evolutionary relationships among the phyl. IIB strains by building a maximum likelihood phylogenetic tree based on the alignment of 3451 genes that are present in single copy in all analyzed genomes. The genome of *R. solanacearum* phyl. IIC strain K60 was used to root the tree. This core genome tree of phyl. IIB is shown in **Figure 2A**. Sequevar assignment, country of origin, and host of isolation were included as well to put the tree into the context of what was already known about these strains.

**Figure 2:**
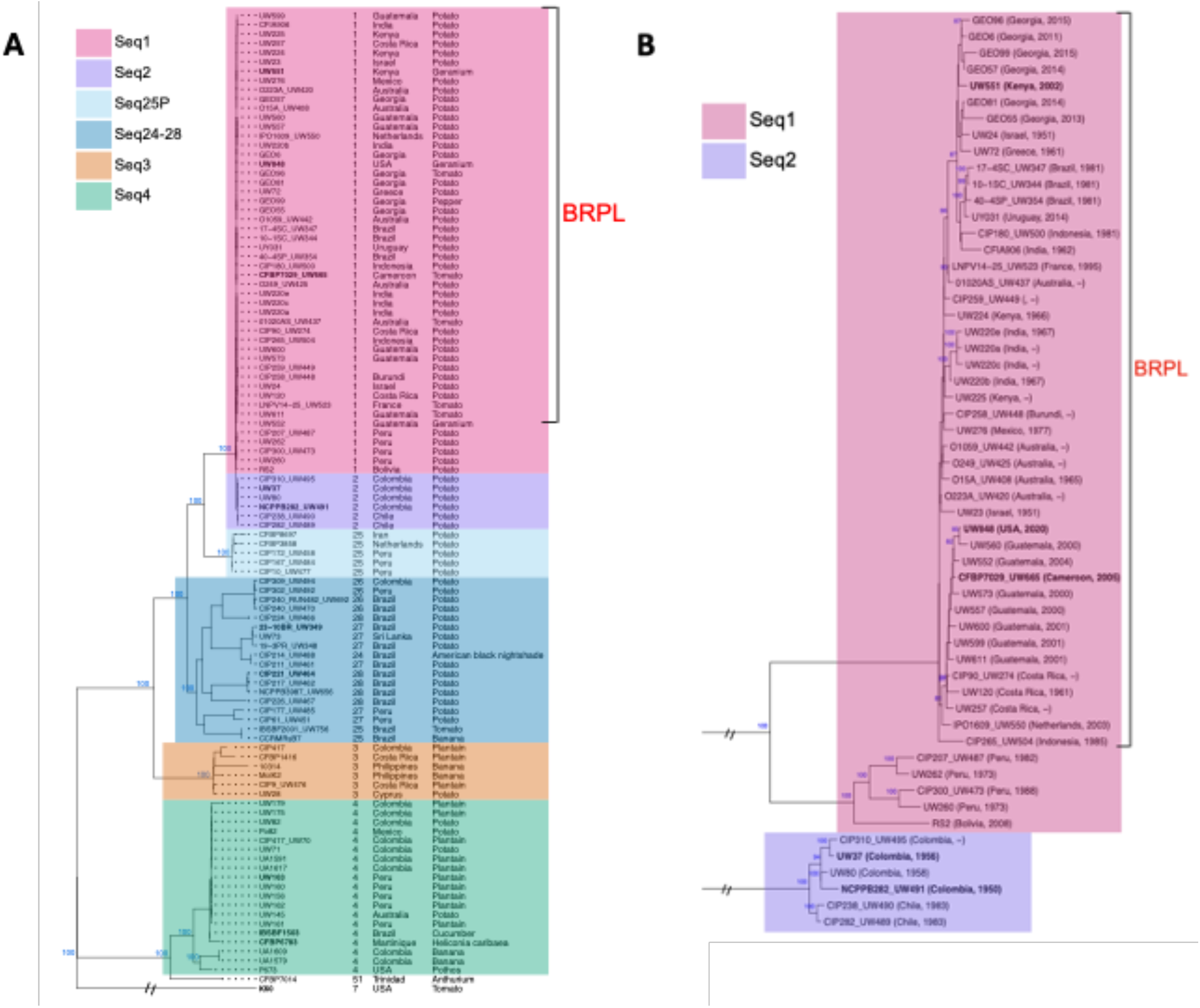
P**h**ylogenetic **analysis of *Ralstonia solanacearum* phylotype (phyl.) IIB. A.** This maximum likelihood phylogenetic tree of 106 phyl. IIB genomes is based on the alignment of 3451 genes and rooted on the genome of phyl. IIC strain K60. Colors represent different sequevars: 1, 2, 3, 4, and 24-28. The brown rot pandemic lineage (BRPL) is labeled. Seq24-28 represents the clade that encompasses strains assigned seq. 24, 26, 27, and 28, as well as some non-Peruvian seq. 25. Seq25P is a clade containing seq. 25 strains, mostly from Peru, that branched separately. The exact sequevar assignment is indicated to the right of each strain name, followed by the country of isolation and the host of isolation. Bootstrap support for all major clades is marked at the nodes of each clade. **B.** Phylogenetic tree using SNP-analysis of only seq. 1 and 2 strains using CIP10_UW477 as the reference. Bootstrap support for high support clades (more than 80%) is marked at the nodes. The country of isolation and the date of isolation are indicated in parentheses for each strain.

The five major clades in phyl. IIB were easily distinguished, each having a bootstrap support of 100. The clade located at the top of the tree correlates perfectly with seq. 1 and 2 and is composed of seq. 1 strains isolated around the world and seq. 2 strains isolated only in South America. Most were isolated from potato, with a few seq. 1 strains from tomato and geranium. Seq. 25 strains isolated from potato in Peru, Iran, and the Netherlands form a clade basal to the node of the seq. 1+2 clade. A third clade is basal to the node of the first two clades and consists of seq. 24, 25, 26, 27, and 28 strains isolated mostly from potato but also from tomato, banana, and American Black Nightshade. Most of these strains originated in Brazil with one strain each from Colombia, Peru and Sri Lanka. The most basal of the top 4 clades consists exclusively of seq. 3 strains. These were isolated in Costa Rica, Colombia, the Philippines, and Cyprus from plantain, banana and (once) from potato. The bottom clade in the tree does not share a most recent common ancestor (MRCA) with the other clades and consists of seq. 4 strains and a single seq. 51 strain. Strains in this clade were isolated from plantain, banana, and potato in North and South America, the Caribbean basin, and Australia. One seq. 4 strain, P673,was isolated from a Pothos plant in Florida, and was previously found to be cool virulent (Bocsanczy et al. 2012), and another, CFBP6783, was from an ornamental plantain (*Heliconia caribaea*) on the Caribbean island of Martinique. The seq. 51 strain was isolated from an Anthurium plant on the Caribbean island of Trinidad.

Correlating the phylogeny with the country of isolation shows that all clades, apart from the top (seq. 1+2) clade, are dominated by strains from South America. This suggests that the MRCA of the strains in each of these clades originated in South America and their progeny were mostly dispersed throughout South America. Phyl. IIB strains were only rarely exported to other continents, with the vast majority of exported strains belonging to seq. 1.

### Phylogeny of *seq. 1 and 2* strains using a reference-based SNP tree

Close inspection reveals substructure within the seq. 1+2 clade (**Figure 2A**). To investigate this clade further, we generated a subtree of the above core genome tree limited to this clade (**Supplemental Figure S1**). This tree revealed that seq. 2 strains form a clade that is distinct from seq. 1. Moreover, five seq. 1 strains form a distinct clade that is basal to all other seq. 1 strains.

To confirm this phylogenetic structure, we built a new tree for only seq. 1 and 2 strains using the genome of phyl. IIB seq. 25 strain CIP10_UW477 (hereafter UW477) as the reference to which reads of all other genomes were aligned (**Figure 2B**). UW477 was chosen since it is the strain with the highest quality genome sequence that is most closely related to strains in seq. 1 and 2. Therefore, we could expect seq. 1 and 2 genomes to align with the UW477 genome over most of their lengths (approximately 5 Mb), allowing us to build a highly resolved tree. This new tree was based on a higher number of informative single nucleotide polymorphisms (SNPs) compared to the SNPs present in the smaller number of genes shared between the phyl. IIB strains and the phyl. IIC strain K60 that was used for the core genome tree (approximately 3.5 Mb).

The most basal clade corresponds to seq. 2 strains of exclusively South American origin (Colombia and Chile) (**Figure 2B**). The second most basal clade consists of seq. 1 strains exclusively of South American origin (Peru and Bolivia). Finally, the most derived clade corresponds to BRPL strains from all over the world, including three strains from Brazil and one strain from Uruguay, but not a single strain from the Andean countries Peru, Bolivia, Colombia, or Chile. Since the two clades basal to the node of the BRPL are all from Andean countries, the MRCA of the BRPL can be inferred to have originated in an Andean country, from which it spread around the world.

A SNP analysis can often give further insights into the time of emergence of a lineage and transmission paths by correlating collection year and geographic location of isolates with their respective position in the phylogenetic tree. However, only 22 SNPs were found between the BRPL isolate with the earliest collection year (UW23, 1951) and the most recent collection year (UW848, 2020), revealing that the mutation rate in the BRPL is extremely low. As a result, there are not enough SNPs to confidently reconstruct transmission paths. In fact, isolates from several countries are found in multiple, distinct positions within the BRPL clade showing that it is generally not possible to infer transmission events between geographic locations. In only one case is the SNP tree consistent with a known route of transmission: the genome of UW848, a BRPL strain isolated from geranium cuttings that were exported from Guatemala to the U.S, is located in a clade for which the genomes of isolates from Guatemala are immediately basal, consistent with the Guatemalan origin of UW848 (Roman-Reyna et al. 2021). Finally, a linear regression analysis for seq. 1 strains showed that year of collection and distance from the MRCA are not correlated (**Supplementary Figure S2**), meaning that it is impossible to infer the time of emergence of the BRPL.

### Pairwise genome similarity of phyl. IIB strains

We previously found that ANI can be used to delineate within-species groups in the RSSC (P. Sharma et al. 2022). We therefore computed pairwise ANI values among phyl. IIB genomes to determine if the clades from the above trees correlate with distinct ANI-based groups. **Figure 3A** shows the resulting heat map, which clearly distinguishes the five major clades of the core genome tree. In addition, seq. 1 and seq. 2 genomes can be distinguished from each other based on ANI. However, ANI, which analyzes only shared (common) gene sequences, cannot distinguish the genomes of the seq. 1 BRPL isolates from the seq. 1 genomes of South American origin because these genomes are intermingled with each other instead of forming separate groups.

**Figure 3:**
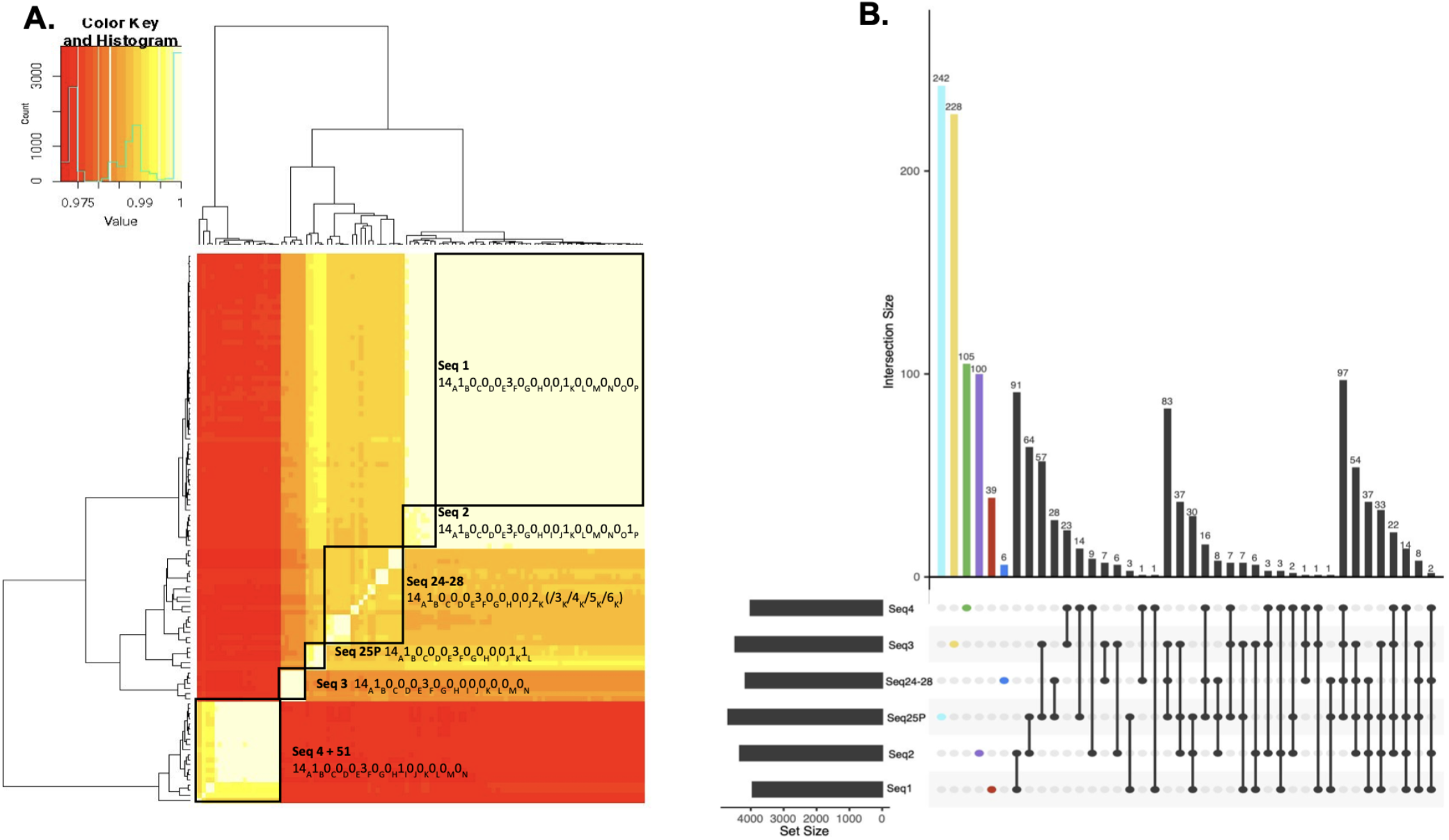
P**a**irwise **average nucleotide identity (ANI) analysis and pangenome analysis of *Ralstonia solanacearum* phylotype (phyl.) IIB.** (A) Heatmap of pairwise ANI percentages between all phyl. IIB strains. Clusters corresponding to each sequevar group from **Figure 2A** are highlighted with their corresponding life identification numbers (LINs). (B) Upset plot representing the pangenome analysis of phyl. IIB strains. Each bar on the chart represents the number of genes present in the corresponding combination of sequevars indicated below the bar. Core-genes for each sequevar group are highlighted in different colors.

To enable genome-based identification of *R. solanacearum* isolates of concern as members of any of the five clades, the corresponding ANI-based groups were circumscribed in the LINbase web server at linbase.org (Tian et al. 2020). The shared component (*i.e.*, the LIN prefix) of the combined seq. 1+2 group is 14_A_1_B_0_C_0_D_0_E_3_F_0_G_0_H_0_I_0_J_1_K_0_L_0_M_0_N_0_O_, corresponding to a minimal pairwise ANI value of 99.9%. The individual seq. 1 and 2 strains are instead circumscribed to the LIN P position, corresponding to a minimum pairwise ANI of 99.925%. LIN prefixes are shown in **Figure 3A**.

### Pangenome analysis and molecular marker design

To investigate differences in gene content between the clades in the core genome tree, we queried the output of the pangenome analysis and identified the core (shared) genes that were unique to individual sequevars and the core genes across sets of sequevars (**Figure 3B, Supplementary Table 2**). For example, 39 genes are exclusively core genes of seq. 1. In other words, while these genes may be present in individual genomes of other seq., they are present in 95% of all genomes of seq. 1. In another example, seq. 1 and seq. 2 genomes have 91 core genes. That is, these 91 genes are present in 95% of all seq. 1 and seq. 2 genomes but are only occasionally present in genomes of other seqs. Overlap in core genes shared among distinct sets of sequevars suggest that the strains in these sequevars acquired and lost similar sets of genes, likely due to adaptation to similar ecological niches.

We were also interested in identifying genes present in all genomes of seq. 1 or seq. 2 or the combined seq. 1+2 clade that were absent from every single genome in every other sequevar, because these genes could be used to develop diagnostic markers and, could potentially provide insights into the biological basis of the distinct fitness of these groups. Five such genes were identified in seq. 1 genomes; that is, they were present in over 95% of seq. 1 genomes but not in a single genome of the other sequevars (**Supplementary Table 2**). One of these genes, a 639-bp gene coding for a hypothetical protein with NCBI gene ID OYQ04669.1 in the UW551 genome, was present in 100% of seq. 1 strains and thus ideally suited for diagnostic marker design (see below). Three of the other four genes also code for hypothetical proteins, while one encodes a predicted HEPN-AbiU2 domain-containing protein with no obvious function in virulence. Thirty-nine genes were unique to seq. 2 strains; that is, these were present in at least five of the six seq. 2 genomes but not in a single genome of the other clades. Of these, 28 code for hypothetical proteins and the other 11 for proteins with predicted functions but, again, none with an obvious involvement in virulence. The combined seq. 1+2 clade had 38 unique genes not present in any genome of any of the other sequevars, of which 21 code for hypothetical proteins and the other 17 have predicted functions without an obvious involvement in virulence. Initial screening suggested that 100% of the BRPL genomes share one unique gene that is absent from all other genomes, a 621-bp gene that codes for a hypothetical protein. However, closer inspection also found the DNA sequence of this gene in the seq. 1 genomes of South American origin. It is not annotated as a gene in these genomes because of a one-nucleotide insertion that caused a frameshift. The presence of this DNA sequence in all seq. 1 genomes therefore precludes its use as a BRPL-specific molecular marker but PCR primers could still be designed flanking the one-nucleotide insertion to distinguish the BRPL from the genomes of South American origin using PCR followed by Sanger sequencing.

We also evaluated the presence of the genes targeted by the primer pairs 630/631(Fegan et al. 1998) and RsSA3 (Fegan et al. 1998; Stulberg et al. 2016). Whereas the gene targeted by RsSA3 is exclusively present in all seq.1+2 strains, the gene targeted by the primers 630/631 (used to identify *Ralstonia* R3bv2 in the European Plant Protection Organization diagnostic protocols) is present in all seq. 1 + 2 genomes and also in the seq. 27 strains UW73, 23-10BR_UW349, and 19-3PR_UW348. These results were confirmed using virtual PCR with the same primer pairs (**Supplementary Table 1**). A third primer set developed to detect R3bv2, the RSCV primers (Stulberg and Huang 2015; Stulberg et al. 2018; Fegan et al. 1998), do not target any gene identified in our analysis. However, virtual PCR showed that this primer pair targets, and is specific to, seq.1+2 genomes, suggesting that the RSCV primers target an intergenic region.

To provide a rapid tool to distinguish between seq. 1 and 2 strains, we developed PCR primers targeting the seq. 1-specific gene (Gene ID: RRSL_RS18155 based on the UW551 genome with accession number GCA_002251655.1). The following primer pair gave a positive virtual PCR result, experimental PCR result, and qPCR result for seq. 1 isolates but not for any genome/strain of any other sequevar (see **Supplementary Figure S3** for details on methods and results): seq1F (CGAATGGACGCCTTACGAGA) and seq1R (CAGGCCGGTCAAAGACTGAA), product size of 161 bp, melting temperature of 60°C.

### Cool virulence phenotyping

To determine if the phylogenomic positions of *R. solanacearum* strains in phyl. IIB could be used to predict their cool virulence, we inoculated plants with a set of ten strains representing the phylogenetic groups identified by the bioinformatic analyses described above. The characteristics of the strains chosen for these assays are detailed in **Figure 1B**. These included three members of the clonal BRPL; South American strains from seq. 2, 4, 27, and 28; and two non-South American controls, the *R. solanacearum* type strain K60^T^ (phyl. IIC seq. 7) and the widely studied model strain GMI1000 (phyl. I seq. 18). Strains were included in phenotypic analyses only if they were sufficiently virulent on a wilt-susceptible cultivar of tomato, which is a nearly universal host of *R. solanacearum*. For this purpose, “virulent” was defined as being able to cause a final mean disease index greater than 2.0 at 28°C, a threshold that required a strain to kill more than half the inoculated tomato plants (**Supplementary Figure S4)**. Establishing this minimal virulence threshold under one condition allowed us to directly compare the tested strains.

The ability to infect and kill plants at low temperatures is believed to make certain *R. solanacearum* strains uniquely threatening. To illustrate this cool virulence phenotype, we characterized two contrasting *R. solanacearum* strains. The representative BRPL strain UW551 and the endemic N. American tomato isolate K60^T^ were indistinguishably lethal on tomato at 28°C (**Figure 4A**). However, these two strains had strikingly different phenotypes on potato at 22°C, with UW551 wilting and killing all inoculated plants, whereas K60 caused no detectable symptoms under these conditions (**Figure 4A**).

**Figure 4.**
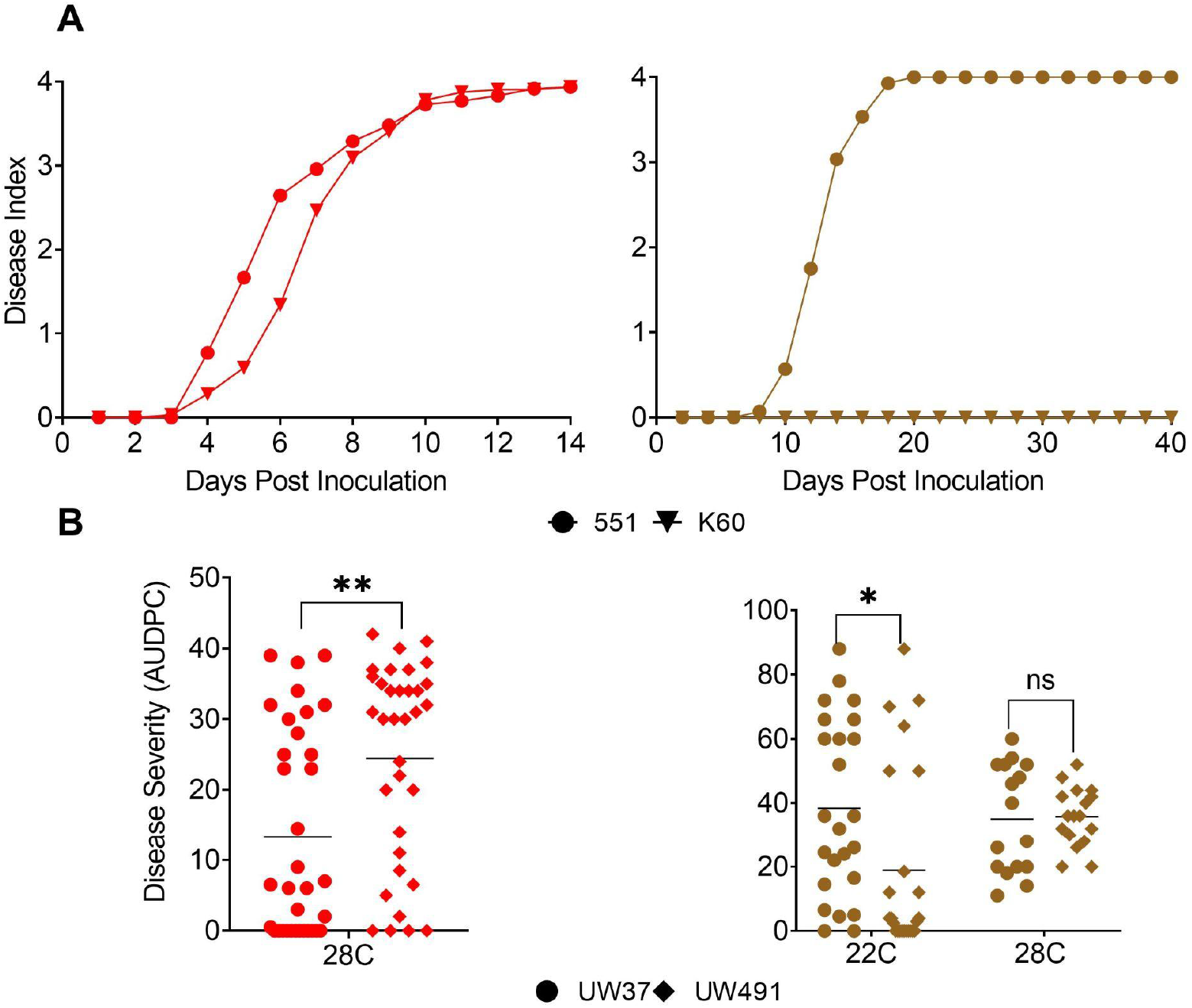
Temperature differentially affects virulence of selected *R. solanacearum* strains on tomato and potato. Unwounded 21-day-old tomato plants (cv. Bonny Best) and 28-day-old potato plants (cv. Russet Norkotah) were inoculated by pouring 50 ml of a 1×10^8^ CFU/ml water suspension of the indicated *Ralstonia* strain into the pot. Plants were incubated in a growth chamber at 28°C (tomato, red symbols) or 22°C (potato, in brown symbols) and rated on a 0-4 disease index scale over 14 days (tomato) or 40 days (potato). **A. Example of the *R. solanacearum* cool virulence phenotype.** Brown Rot Pandemic Lineage (BRPL) strain UW551 and N. American strain K60 had similar virulence on tomato (in red) at 28°C (*P*=0.104, Two-way Repeated Measures ANOVA), but at 22°C only UW551 was virulent on potato (in brown) (*P*<.0001, Two-way Repeated Measures ANOVA). Each data point indicates the mean disease index of 18 to 48 replicate plants in 3 to 8 independent experiments. **B. Host preference and cool virulence can interact.** *R. solanacearum* seq. 2 strain UW491 was more virulent than seq. 2 strain UW37 on tomato plants (red symbols) at 28°C (*P*=0.0022, Welch’s t test), but not on potato plants at 28°C (brown symbols) at 28°C (*P*=0.995, Šidák’s post hoc test). Conversely, UW37 was more virulent than UW491 on potato at 22°C (*P*=0.010, Šidák’s post hoc test). Each symbol indicates the Area Under the Disease Progress Curve (AUDPC) for a single plant; data for each condition represent 16 to 34 replicate plants, in 2 to 5 independent experiments.

The historical literature assumes that cool virulence is a qualitative trait that clearly distinguishes uniquely threatening *Ralstonia* strains. To explore this assumption, we measured the virulence of representative strains from various clades on wilt-susceptible tomato at 28°C and 22°C (**Supplemental Figure S5A**). We defined cool virulence as the ability to reach a mean disease index >2.0 on tomato plants at 22°C, which corresponds to killing more than half the inoculated plants.

Using this definition, strains UW551, K60, and CFBP6783 are cool virulent on tomato. Phyl. IIB-1 strain UW551, a member of the BRPL isolated from geranium, was the most virulent of all strains on tomato at 22°C. Strains K60 (phyl. IIC-7, from tomato) and CFBP6783 (phyl. IIB-4, from *Heliconia caribaea*) exhibited moderate cool virulence, the model *Ralstonia* strain GMI1000 (phyl. I-18, from tomato) had low cool virulence, and strain UW349 (phyl. IIB-27, from potato, classified as R3bv2 by EPPO) was not cool virulent, causing no symptoms on tomato at 22°C. These data demonstrate that, on tomato, cool virulence is a quantitative trait. Given that all these strains caused symptoms at 28°C, we conclude that the ability of a *Ralstonia* strain to cause disease at 28°C does not predict its virulence on the same host at 22°C.

As mentioned above, most *Ralstonia* strains can infect and cause wilt symptoms on both the fast-growing and easily assayed tomato, and potato, the more experimentally challenging host of primary regulatory interest. When we compared the ability of individual strains to infect both plants, we found that strain, host, and temperature interacted in unpredictable ways. For example, we compared the virulence on tomato and potato of UW37 and UW491, which are phyl. IIB-seq. 2 strains isolated from potato in Colombia. Based on core genome phylogeny, these strains are nearly identical (**Figure 2**). At 22°C on potato, strain UW37 caused significantly more disease than UW491 (**Figure 4B**). To determine if this phenotypic difference persists at 28°C, we inoculated potatoes at 28°C and found that at a warmer temperature they caused similar levels of disease (**Figure 4B**). Unexpectedly, when we inoculated tomato plants at 28°C with these strains, the outcome was reversed with strain UW37 causing significantly less disease than UW491 (**Figure 4B**). In summary, the only reliable predictor of these strains’ cool virulence on the primary host of regulatory interest was a direct phenotypic assay on potato plants.

Because potato plants suffered abiotic (heat) stress when incubated at 28°C, subsequent potato inoculation assays were conducted only at the production-relevant temperature of 22°C. These assays measured the AUDPC of individual strains following soil-soak inoculation of unwounded Russet Norkotah potato plants (**Figure 5)**. We observed a nearly continuous distribution of cool virulence levels across the strains tested. As expected, phyl. IIC-7 strain K60 caused no symptoms under these conditions. The three BRPL strains, UW551, UW848, and UW665, were the most virulent. Two seq. 2 strains, UW37 and UW491, which are also regulated as R3bv2, were only moderately virulent on potato at 22°C. In contrast, seq. 28 strain UW464, which belongs to a distinct clade and is not regulated as R3bv2, was as cool virulent as BRPL strain UW665. The tropical lowland seq. 4 strains UW163 and CFBP6783 did not differ in cool virulence on potato from the seq. 2 strain UW37, nor from phyl. I-18 model strain GMI1000, which was originally intended as a non-cool virulent control. Together, the results of these assays demonstrate that cool virulence is a quantitative trait on potato as it is on tomato. Moreover, these results show that cool virulence is not limited to currently regulated R3bv2 strains.

**Figure 5.**
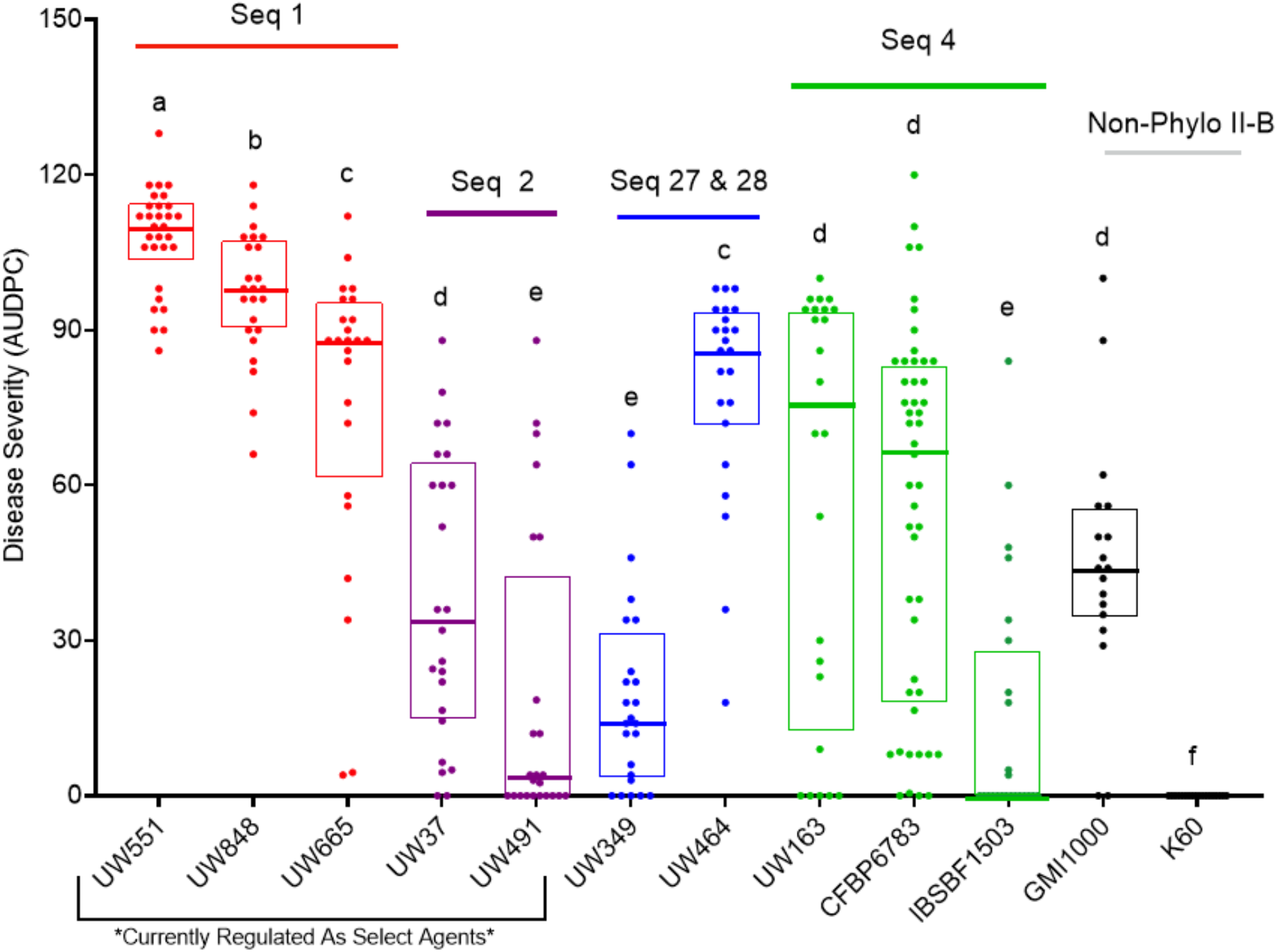
Cool virulence on potatoes is not limited to *Ralstonia* strains designated as Race 3 Biovar 2 strains, nor to those designated as Brown Rot Pandemic Lineage strains. Four-week-old Russet Norkotah potato plants growing at 22°C were inoculated by soil drench with 1 x 10^8^ CFU of the indicated *Ralstonia* strain and rated visually for wilt symptoms over 40 days. Each symbol indicates the Area Under the Disease Progress Curve (AUDPC) for a single plant. Horizontal bars indicate the median AUDPC for each strain; data shown are from three to eight independent experiments, each containing 6 plants/strain. Strains labeled with the same lower-case letters have indistinguishable virulence (*P*>0.05, Repeated Measures ANOVA).

### Colonization of potato crowns and daughter tubers

Having determined that multiple strains not designated as R3bv2 can wilt potato plants at cool temperatures, we hypothesized that the global success of the BRPL derives not from a unique ability to cause symptoms but rather from a differential capacity to latently infect and colonize potato plants and tubers under cool conditions. We tested this hypothesis by measuring the population sizes of a subset of *Ralstonia* strains in potato plant crowns and tubers after natural infection through unwounded roots. After two to three weeks, plants were classified as symptomatic or asymptomatic based on visible wilt symptoms. Tissue samples from the crown (stem just above soil level) and two tubers from each plant were ground and serially dilution plated to determine the CFU/g tissue. **Figure 6** shows the *Ralstonia* population sizes in stems and **Figure 7A** shows bacterial population sizes in tubers. In these figures, data points from symptomatic and asymptomatic plants are distinguished as closed circles and open squares, respectively.

**Figure 6.**
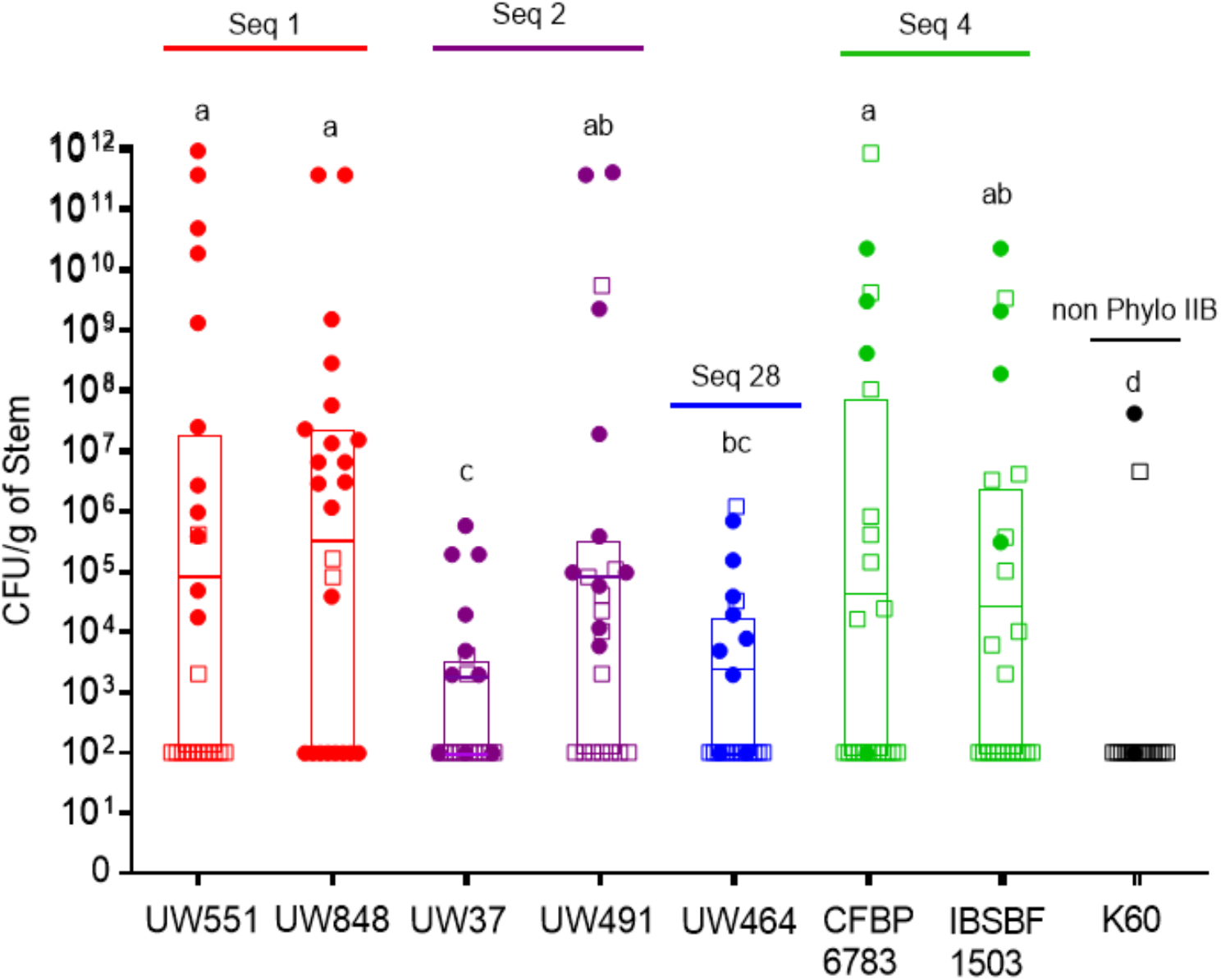
Diverse phylotype IIB *Ralstonia* strains effectively colonize potato stems. Population sizes of diverse *Ralstonia* strains in stems that were harvested 2-3 weeks after potato plants were inoculated by soil drench and incubated at 22°C. A 0.1-g stem slice from each plant was surface sterilized, ground in water and serially dilution plated on SMSA medium. Samples with bacterial burden below the detection limit (2×10^2^ CFU/g) were plotted at 10^2^ CFU/g. Samples harvested from asymptomatic plants are indicated by open squares; samples harvested from symptomatic plants are indicated by filled circles. Horizontal bars indicate the geometric mean population sizes for each experimental status (symptomatic vs. asymptomatic plants). The frequency of samples with undetectable populations for this experiment is shown in Supplementary Fig 5. One-way ANOVA using multiple Student’s *t*-test comparisons was performed on pooled data from all symptomatic and latently infected stems that contained detectable populations of *Ralstonia*. No strain by status interaction was observed. Data for each strain are from 17 to 24 replicate plants in 3 to 4 independent experiments.

**Figure 7.**
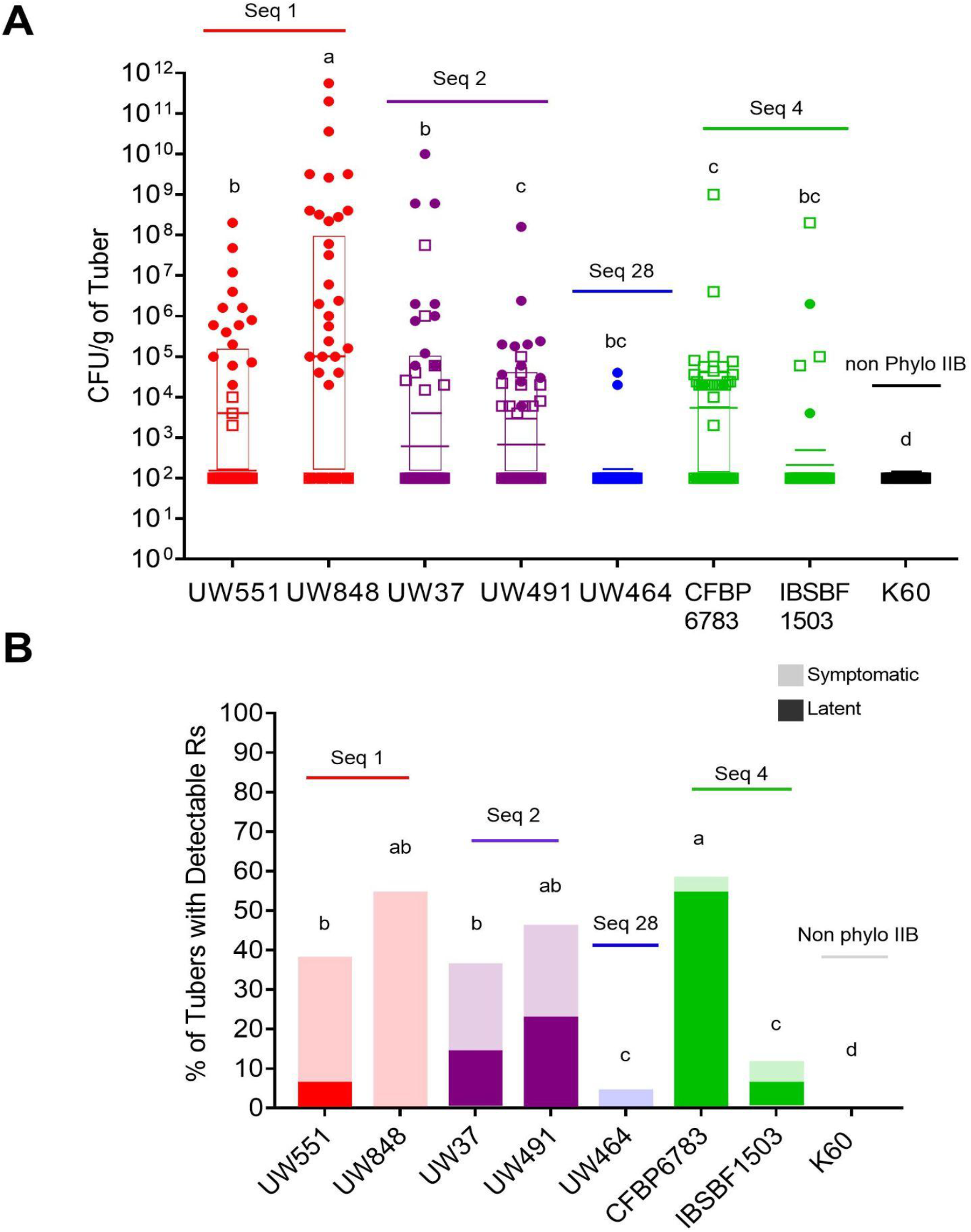
Cool virulence of *Ralstonia* strains on potato does not predict ability to form latent infections or colonize potato tubers. A. Population sizes of diverse *Ralstonia* strains in cv. Russet Norkotah tubers that were harvested 2-3 weeks after plants were inoculated by soil drench and incubated at 22°C. The 2 largest healthy-looking tubers from each plant were surface sterilized and a 0.1-g core from the stolon end was ground in water and serially dilution plated on SMSA medium (detection limit= 2×10^2^ CFU/g; samples without detectable *Ralstonia* cells were assigned a value of 10^2^ CFU/g). Samples derived from asymptomatic plants are indicated by open squares; samples derived from symptomatic plants indicated by closed circles. Geometric mean population sizes are indicated by horizontal bars. Strains labeled with different lower-case letters have different population sizes in tubers (*P*<0.05, One-Way ANOVA with multiple student’s t-test comparisons). Statistical analyses were performed on pooled symptomatic and latently infected tubers with detectable populations. No interaction between strain and symptomatic status was observed. Data for each strain include 34 to 48 replicate samples from 3 to 4 independent experiments. B. Frequency of detectable *Ralstonia* cells in potato tubers. Each bar indicates the percentage of tubers with detectable *Ralstonia* populations from plants that were asymptomatic (darker lower bar) and symptomatic (lighter upper bar) for each strain. Lower case letters over each column indicate differences among strains (*P*<.0001, Pearson chi square test); multiple statistical comparisons between strains were calculated using the odds ratios post hoc test.

All tested *Ralstonia* strains were detected in at least some sampled stems, with many samples containing more than 1×10^4^ CFU/g tissue (**Figure 6**). Bacterial population sizes in stems varied broadly within strains, consistent with the stochastic nature of root infections. Five of the eight tested strains fell into a statistically indistinguishable cluster that all reached a mean population size >5×10^4^ CFU/g, based on combined data from symptomatic and asymptomatic plants. Unsurprisingly, populations of most strains tended to be higher in samples from symptomatic plants. The two tested BRPL strains, UW551 and UW848, reached the highest mean population sizes during symptomatic infection. We defined latently infected plants as those with detectable populations (here > 10^3^ CFU/g) but no visible symptoms. On this basis, the BRPL strains rarely caused latent infections under these conditions. In contrast, seq. 4 strains often caused latent infections, and one of the strains, CFBP6783, reached the largest population size detected for any strain, and this was in an asymptomatic plant. Phyl. IIB-2 strain UW491 colonized stems as well as BRPL and phyl. IIB-4 strains, reaching populations significantly higher than those of the closely related seq. 2 strain UW37 (**Figure 6**). This was unexpected because UW37 was more virulent on potato than UW491 under these conditions (**Figure 4B**). Despite having cool virulence indistinguishable from that of a BRPL strain, IIB-28 strain UW464 colonized potato stems relatively poorly. Although negative control strain K60 never caused disease in soil soak experiments, it did colonize the stems of 2 of 17 plants tested.

Importantly, many stem samples contained no detectable *Ralstonia* cells (the detection limit of the dilution plating was 2×10^2^ CFU/g). A qualitative analysis of detection frequency found that apart from K60 all strains were similarly likely to reach detectable thresholds in stems (**Supplementary Figure S6**). Together, these results indicate that potato stem colonization is a complex behavior that is not unique to the BRPL.

Ability to colonize daughter tubers is epidemiologically important because brown rot is often disseminated when latently infected tubers are planted as seed. We measured population sizes of *Ralstonia* strains inside surface-sterilized daughter tubers formed by inoculated potato plants. Any tubers with visible rotting were not sampled. As for stems, in tubers we observed substantial within-strain variability in *Ralstonia* population sizes, with generally higher populations from symptomatic than asymptomatic plants (**Figure 7A)**. The BRPL strains often reached tuber populations >10^5^ CFU/g, but seq. 2 strains, which are also regulated as R3bv2, trended towards lower population sizes (**Figure 7A**). The strains that had the smallest population sizes in stems, UW464 and K60, also had the lowest tuber population sizes. In contrast, CFBP6783, a phyl. IIB-4 strain that reached high populations in stem tissue, had relatively low populations in tubers. These results suggest that stem colonization is not a reliable predictor of tuber colonization and that tuber colonization, like stem colonization, is not a reliable predictor of the ability to wilt potato plants.

In contrast to the results from stem tissue, *Ralstonia* strains varied substantially in the frequency with which they colonized tubers (**Figure 7B**). Most strains were detectable in fewer than half the sampled tubers. Three tested strains were rarely (UW464, IBSBF1503) or never (K60) detected in tubers. Only BRPL strain UW848 and phyl. IIB-4 strain CFBP6783 colonized more than 50% of the tubers at the time of sampling, 2 to 3 weeks after inoculation. However, while all the tubers that tested positive for UW848 came from symptomatic plants, a large majority of the tubers that contained detectable cells of CFBP6783 were from symptomless plants (**Figure 7B**).

In summary, the results of the colonization assays show that the ability to infect, persist, and multiply in potato stems and tubers at cool temperatures is not unique to R3bv2. Although strains in the BRPL were among the best colonizers of potato tissue, this trait is shared with closely related seq. 2 strains and also with geographically and phylogenetically distant strains in seqs. 28 and 4.

## Discussion

*R. solanacearum* causes one of the most destructive diseases of potato, potato bacterial wilt or brown rot. Concerns regarding the introduction of exotic cool virulent strains into northern regions have motivated regulations to restrict the introduction and movement of R3bv2 strains. The evolutionary and genetic basis of the cool virulence phenotype is also of fundamental interest. Therefore, here we integrated a comparative evolutionary genomics analysis with biologically relevant phenotyping to better understand the nature and distribution of the cool virulence phenotype among *R. solanacearum* strains.

A previous genome-enabled phylogeographic analysis of 17 phyl. IIB isolates suggested that most potato brown rot outbreaks outside of South America were due to a single clonal lineage that probably originated in South America (Clarke et al. 2015). We designated this lineage the Brown Rot Pandemic Lineage (BRPL). The previous phylogeographic analysis was limited by the relatively low quality of genome assemblies (which allowed the use of only 2 Mb of each genome for phylogenetic reconstruction) and the small number of genomes of both South American origin and from around the world (which limited the confidence in the phylogeographic reconstruction). Here, we used 106 high quality phyl. IIB genomes in our phylogeographic analysis. This significantly improved the phylogenetic reconstruction and identified the likely geographic origin of the BRPL. In particular, a SNP analysis using the entire length of the 56 genomes of the BRPL isolates and the closest BRPL relatives in seqs. 1 and 2 provided a highly resolved phylogenetic tree. In this tree, all 11 isolates on the two branches basal to the inferred MRCA of the 45 BRPL isolates were collected in the Andean highlands of Peru and Bolivia and neighboring regions in Chile and Colombia. Therefore, we can now conclude with high confidence that the BRPL emerged in the Andean highlands, the center of potato genetic diversity.

Other phyl. IIB potato-infecting strains also likely emerged in the Andean highlands but, in contrast to the BRPL, these have rarely spread beyond this region. Only two brown rot isolates outside of the BRPL clade were found in countries outside the Andean region: the closely related seq. 25 isolates CFBP8687 from Iran and CFPB3858 from the Netherlands. Both of these strains cluster in a clade with Peruvian isolates, suggesting that they are of Andean origin. Seq. 25 strains have not detectably spread beyond the Netherlands and Iran. Also, one seq. 2 strain reported in Kenya (K. Sharma et al. 2022) does not appear to have spread further. Thus, BRPL clade strains are distinct from other clades in their widespread global distribution.

We had hoped to also infer when the BRPL may have emerged and along which routes it may have been transmitted around the world. However, the extremely low mutation rate in the BRPL and the lack of correlation between the time from the MRCA and the distance from the MRCA of the BRPL isolates made this impossible. In fact, although our analysis included isolates collected over 69 years, the earliest and latest isolates were separated by only 22 SNPs over the entire length of their genomes. This result is consistent with an analysis of BRPL isolates collected over many years from various locations in the Thames River in the UK, whose genomes were also nearly identical apart from minor phenotypic variation likely due to variation in the accessory genome (Parkinson et al. 2013; Farnham et al. 2022).

One explanation for the extremely low mutation rate in the BRPL, and in the RSSC more broadly, could be that these pathogens replicate only slowly in soil and water, experiencing long periods of stasis. Although these periods are punctuated with short bursts of growth in the xylem when the strains infect plants and reach high population densities, the xylem is not known to be a significant inoculum source for brown rot outbreaks. Moreover, long-distance movement in latently infected tubers is relatively rare and our experimental data show fairly low bacterial population sizes in asymptomatic tubers. This scenario is quite different from most other plant and animal pathogens, for which transmission mostly occurs after exponential growth in a host, giving ample opportunity for mutation and differential selection of pathogens prior to transmission.

Why did the BRPL succeed in its worldwide conquest while its close relatives did not? This success is often attributed to cool virulence. However, much evidence indicates that cool virulence is not unique to the R3bv2 group. Cool virulence was previously observed in seq. 4 (Bocsanczy et al. 2012; Norman et al. 2009; Bocsanczy, Huguet-Tapia, and Norman 2017) and this is confirmed by our experiments using different seq. 4 strains. Moreover, although seq. 2 strains are currently strictly regulated in North America and Europe, no seq. 2 strains have been rigorously tested for cool virulence until now.

We wondered if strains currently defined as R3bv2, including those in seq. 2, are universally cool virulent, and if strains outside of R3bv2 beyond seq. 4 also have this trait. We therefore characterized the cool virulence of a set of diverse *R. solanacearum* phyl. IIB strains. We chose representative strains from within seqs. 1 and 2, from other phyl. IIB strains isolated from potato in Andean countries, and from seq. 4 strains from other hosts in the hot lowland tropics. The choice of strains for phenotyping was limited by both the paucity of seqs. 1 and 2 strains isolated from Andean countries and the unfortunate loss of virulence in many culture collections of *R. solanacearum* strains. Additional strains in these groups should be isolated and phenotyped.

Our measurements of the cool virulence of these strains on potato demonstrated that this phenotype is quantitative rather than qualitative. All of the seq. 1 and 2 strains tested were able to infect and wilt at least some potato plants at 22°C (summarized in **Table 2**), suggesting that this qualitative ability is a universal trait within these sequevars. However, the strains varied in their virulence at 22°C, with the seq. 1 strains exhibiting much greater degree of cool virulence than the seq. 2 strains. Importantly, we also found that cool virulence on potato is present in multiple genetic lineages within phyl. IIB. Thus, this trait is not unique to the sequevars within R3bv2 or seq. 4.

**Table 2.**
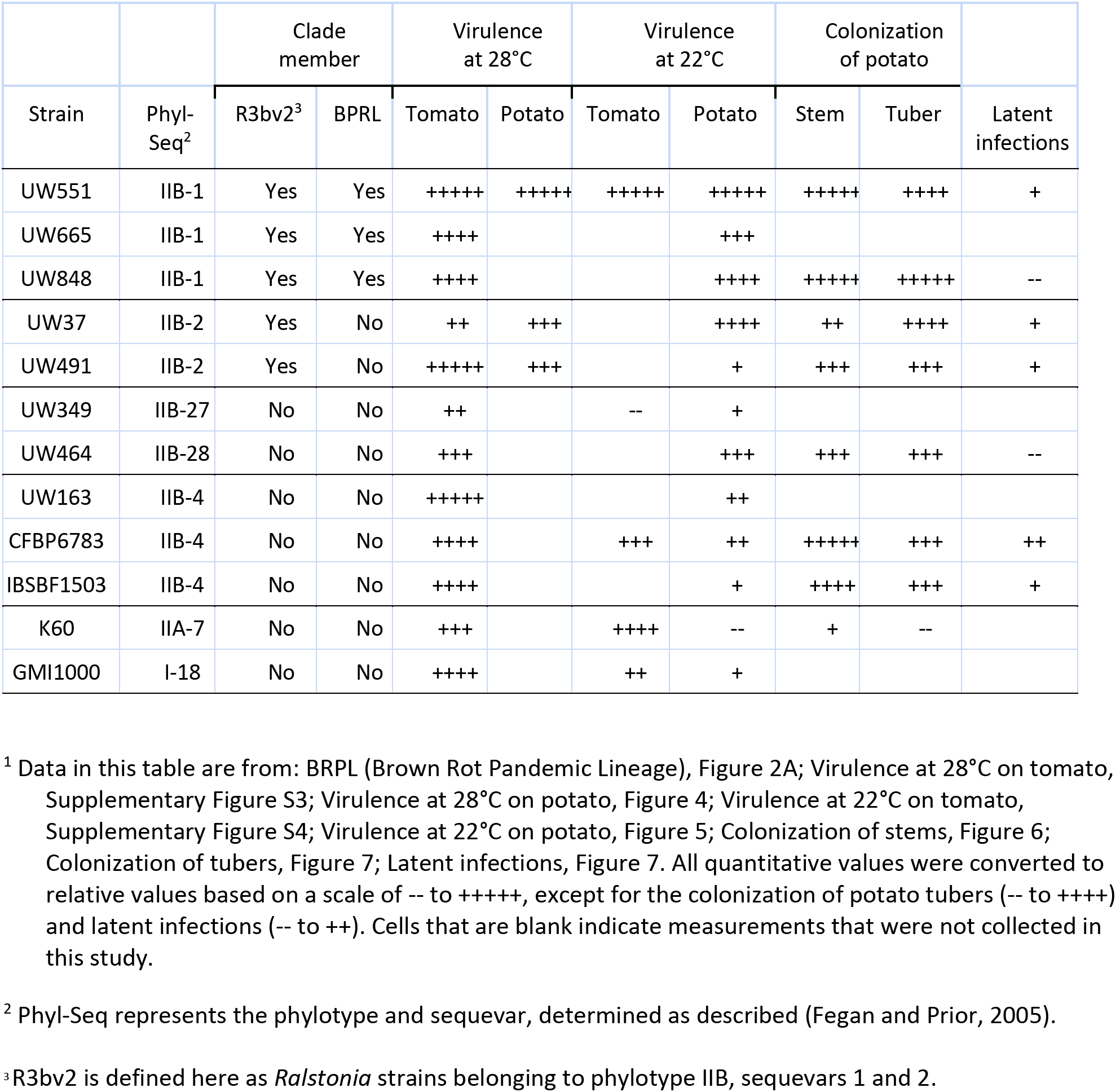
Summary of genomic and phenotypic characterization of selected *Ralstonia solanacearum* species complex strains in this study^1^.

Measuring the cool virulence of isolates on potato is a time-consuming process, requiring as long as 40 days to adequately assess symptoms. With the goal of reducing this time, we asked if the fast-growing model species tomato could serve as a proxy for potato. It takes only 21 days to assess wilt disease on tomato at 22°C. However, a comparison of the cool virulence of the tested *Ralstonia* strains on tomato versus potato found congruence between the two hosts for some strains, but not for all. For example, the closely related seq. 4 strains CFBP6783 and UW163 were both moderately cool virulent on potato, but only UW163 was cool virulent on tomato (data not shown for CFBP6783). Similarly, strains UW349 (seq. 27) and UW464 (seq. 28) had cool virulence on potato, but neither strain was cool virulent on tomato (data not shown for UW464). In contrast, North American phyl. IIA-7 strain K60 was moderately cool virulent on tomato, but was completely unable to cause disease on potato at 22°C. Finally, the closely related IIB-2 strains UW37 and UW491 had opposite patterns of virulence on tomato and potato at high and low temperatures. We conclude that cool virulence on tomato is not a reliable predictor of a strain’s behavior on potato.

Furthermore, a strain’s cool virulence on potato is not reliably predicted by that strain’s phylogenetic relatedness to other strains, its virulence on tomato at a warm temperature (28°C), or even by its virulence on potato at 28°C. This complex pattern suggests that host– and temperature-dependent factors that influence virulence vary not only among phylotypes, sequevars, and other clades within the RSSC, but can also vary within clades of high genetic similarity, such as seq. 1, as previously described by Cellier and Prior (2010) and Wang et al. (2019). Clues to these variations may be present in strains’ accessory genomes. The possibility that some seq. 1 strains tested in previous papers had lost virulence in storage cannot be ruled out.

Given that strains outside of the BRPL exhibit cool virulence, we hypothesized that the global success of the BRPL derives not from cool virulence per se but rather from the ability to form latent infections in tubers under cool conditions, as latent infections would allow these strains to spread undetected during local, regional and global trade. Thus, we evaluated whether the ability to form latent infections and colonize potato stems and tubers under cool conditions was unique to BRPL strains. Among the seven selected phyl. IIB strains tested, the two BRPL strains aggressively colonized both stems and tubers at 22°C, as expected. The two tested seq. 2 strains were actually more likely than the seq. 1 strains to induce latent rather than symptomatic infections at 22°C and also often reached high populations in tubers. Importantly, the two seq. 4 strains, which exhibited moderate to high cool virulence on potato, also induced latent infections and reached high population sizes in potato tubers. This latter result is consistent with the finding that seq. 4 strain P673, isolated from Pothos in Florida, is as virulent on potato as BRPL strain UW551 at the very low temperature of 18°C (Bocsanczy et al. 2012). *R. solanacearum* seq. 4 strains have been isolated in the warm lowland tropics around the Caribbean from dicotyledonous hosts like tomato and cucurbits, and also from monocots like banana, Heliconia, and Pothos. Some seq. 4 strains have curiously specific host ranges, such as IBSBF1503 and UW163, which are respectively pathogenic on melons but not bananas and vice-versa (Ailloud et al. 2015). The fact that multiple seq. 4 strains can colonize potato tubers as well as strains in the BRPL is cause for concern because potato brown rot is often disseminated in latently infected seed tubers.

Our results and those of some previous studies cited below indicate that cool virulence on potatoes is a quantitative trait that cannot be explained by one or a few genes. A genome-wide association study (GWAS) could be used to identify genes or gene alleles contributing to this trait, taking advantage of the genomes sequenced here to go beyond previous studies that have already proposed some candidate genes involved in cool virulence (Norman et al. 2009; Bocsanczy, Huguet-Tapia, and Norman 2017); (Stulberg and Huang 2015; Stulberg et al. 2018; Fegan et al. 1998). However, many more isolates will need to be phenotyped to obtain the necessary statistical power. This is a daunting task since cool virulence assays on potatoes are labor intensive and seq. 1 and 2 strains can only be phenotyped in N. America and Europe under restrictive high-containment conditions.

Our genomic analyses of potato brown rot strains in the BRPL from around the world did not identify candidate genes that could explain the extraordinary fitness of this near-clonal subgroup of seq. 1. Our phenotyping studies likewise found that, under our experimental conditions, the ability to infect and wilt potato plants at cool temperatures is not unique to the BRPL or to currently defined R3bv2 strains. We suspect that our experiments did not capture the particular environmental or biological conditions in which BRPL strains succeed while other *Ralstonia* strains do not. Using consistent inoculum levels, plant genotypes, temperature, and light to study plant-bacterial interactions yields replicable results in manageable time-frames. However, such reductionist experiments do not replicate the environment of a brown rot outbreak in the field. The BRPL has thrived and spread around the world in the presence of diverse abiotic stresses, other pathogens, and agricultural microbiomes, to name only a few variables. A comparison of the virulence of BRPL and seq. 4 strains on potato plants grown under typical field conditions in the highland tropics is needed to verify if their cool virulence in the lab truly reflects their cool virulence as expressed in the field.

How could the results reported here inform the regulation of cool virulent strains? On one hand, our finding that cool virulence is a quantitative trait makes it challenging to clearly distinguish between strains that threaten potato production in northern potato-growing regions and those that do not. On the other hand, the highest level of cool virulence among the phyl. IIB clade strains examined was restricted to seq. 1 while the tested seq. 2 strains had cool virulence similar or lower than seq. 4 strains already established in the United States. Considering these results in the context of the other available experimental and epidemiological data, the BRPL appears to possess a unique set of traits that make it singularly capable of persisting and infecting potato at cool temperatures. Unfortunately, no fully virulent seq. 1 strains of South American origin were available for phenotyping, preventing us from determining their level of cool virulence. BPRL strains could not be distinguished from the seq. 1 strains of South American origin based on gene content and ANI. Therefore, separating the BRPL from seq. 1 of South American origin from a regulatory perspective is impractical with existing data.

Alternatively, the R3bv2 definition could be restricted to seq. 1. In fact, seq. 1 and seq. 2 genomes can be assigned to separate ANI-based LINgroups. Moreover, we identified a gene exclusively present in all seq. 1 genomes. This finding allowed us to design and validate a primer pair specific to seq. 1 that would offer a practical diagnostic test. Although using the designation ‘seq. 1’ for R3bv2 is imperfect because sequevars are defined by the allele sequence of a single gene, the *egl* gene, the seq. classification system is well established in the literature and is supported by extensive data (Cellier et al. 2023). Therefore, “*Ralstonia solanacearum* seq. 1” could be used as a name for a revised R3bv2 definition. Diagnostic tests for this revised R3bv2 could include whole genome-based identification based on ANI, with a clear LINgroup identified using the LINbase database (Tian et al. 2020), PCR based on the gene we found to be exclusively present in seq. 1, and the *egl* allele sequence of seq. 1.

In conclusion, combining thorough genomic and phenotypic analyses can help build the foundation to circumscribe truly threatening pathogenic agents. Such circumscription is critical to minimize both under-regulation that could harm agriculture and over-regulation that could unnecessarily burden scientific research and agricultural commerce. However, in the case of cool virulence in *R. solanacaerum*, a complex genetic basis of a quantitative phenotype requires careful evaluation of all available genetic, phenotypic, and epidemiological data when delimiting the group of organisms to be regulated.

## Abbreviations

RSSC, *Ralstonia solanacearum* species complex; phyl., phylotype; seq., sequevar; BRPL, brown rot pandemic lineage; R3bv2, Race 3 biovar 2; MRCA, most recent common ancestor; AUDPC, Area Under the Disease Progress Curve; LIN, Life Identification Number.

## Acknowledgements

This material was made possible, in part, by a PPA7721-funded Cooperative Agreement from the United States Department of Agriculture’s Animal and Plant Health Inspection Service (APHIS). It may not necessarily express APHIS’ views. This research was supported in part by the U.S. Department of Agriculture (Hatch # WIS04091 to C. Allen; Hatch #1023861 to T. Lowe-Power; NIFA Award #2022-67013-36272 to T. Lowe-Power) and by the University of Wisconsin-Madison College of Agricultural and Life Sciences

