## Supplemental Tables and Figures for "Genotypic and Phenotypic Analyses Show *Ralstonia solanacearum* Cool Virulence is a Quantitative Trait Not Restricted to “Race 3 biovar 2”"

**Supplementary Tables and Figures**

Dewberry RJ, Sharma P, et al**.** Genotypic and Phenotypic Analyses Show *Ralstonia solanacearum* Cool Virulence is a Quantitative Trait Not Restricted to “Race 3 biovar 2”

**Supplementary Table S1. The 106 *Ralstonia solanacearum* phyl. II strains used in this study: Strain name, biological metadata, virtual PCR results, and genome sequence characteristics.**
Dewberry et al Supplementary Table S1.xlsx https://osf.io/3c2fx

**Supplementary Table S2. List of core genes for *Ralstonia solanacearum* phyl. IIB strain in seq1; seq2; seq1+2; and Brown Rot Pandemic Lineage (BRPL), along with genes exclusive to each group.**
Dewberry et al Supplementary Table S2.xlsx. https://osf.io/c7ydb

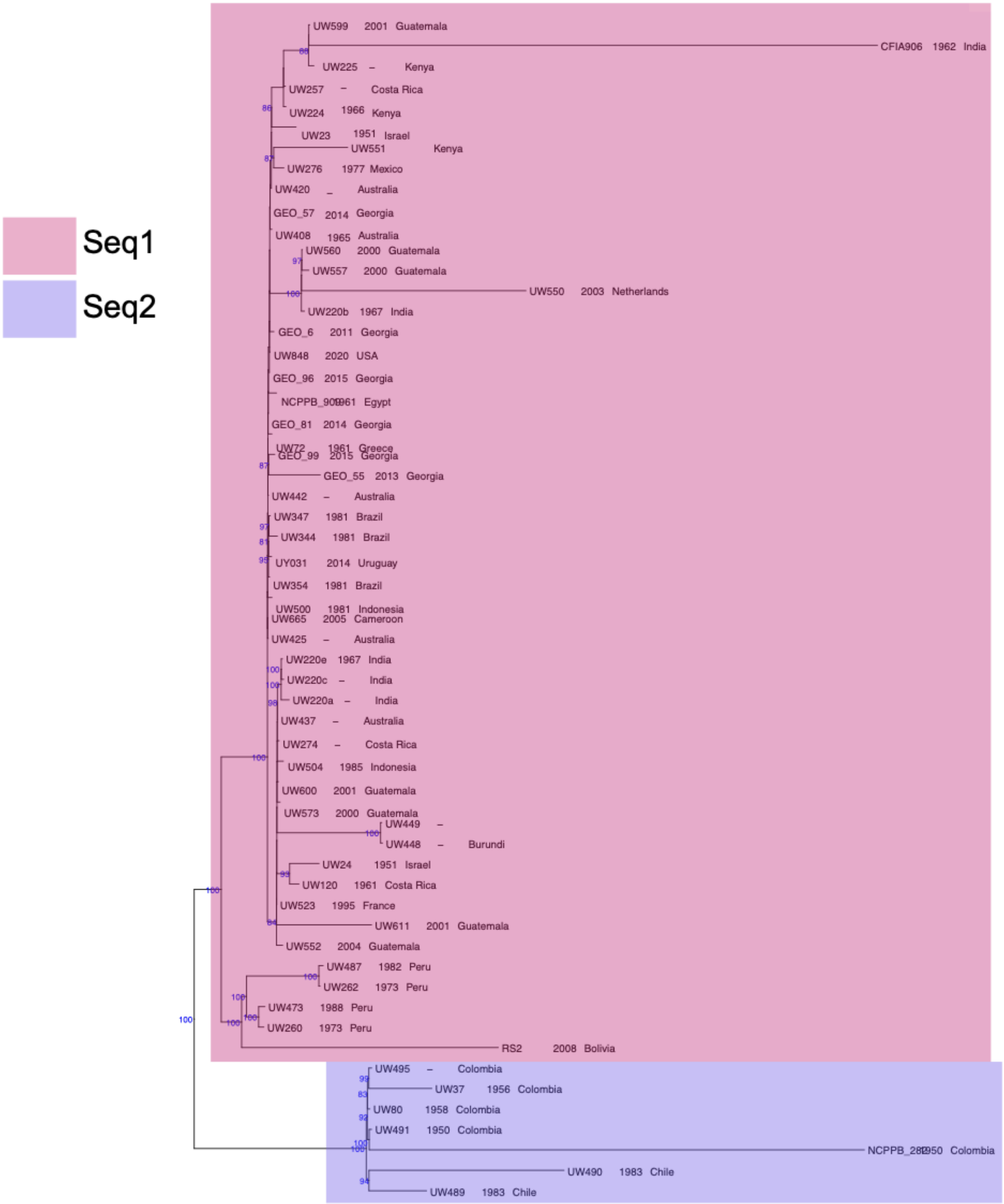


**Supplementary Figure S1: Phylogenetic analysis of only seq. 1 and 2 strains using core-genome analysis**. The maximum-likelihood phylogenetic tree shown in **Figure 1A** was reduced to seq. 1 and 2 genomes. Bootstrap support for high support clades (more than 80%) is marked at the nodes. Sequevar assignments for the strains are highlighted in colors representing seq. 1 and 2. The country of isolation and the date of isolation are indicated next to each strain name.

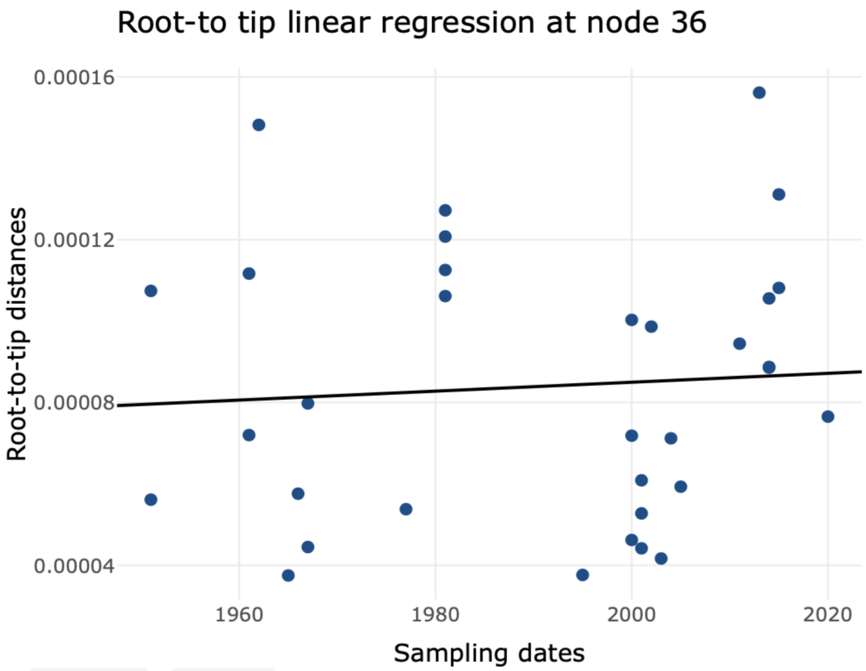


y = −0.00013 + 1.1e−07 x
p-value = 0.696
adjusted $R^2$ = −0.027
x-intercept = 1181.82

**Supplementary Figure S2: Root-to-tip regression analysis of the genomes of brown rot pandemic lineage (BRPL) isolates.** Screenshot from Phylostems (Doizy et al. 2023). A temporal signal analysis was used to evaluate molecular-tip dating for the BRPL genomes illustrated through linear regression analysis of evolutionary distance over sampling dates.

**A**

Primer pair Seq1 (Tm 60C)

| | Sequence (5'->3') | Template strand | Length | Start | Stop | Tm | GC% | Self complementarity | Self 3' complementarity |
|---|---|---|---|---|---|---|---|---|---|
| Seq1F | CGAATGGACGCCTTACGAGA | Plus | 20 | 414 | 433 | 59.90 | 55.00 | 4.00 | 0.00 |
| Seq1R | CAGGCCGGTCAAAGACTGAA | Minus | 20 | 574 | 555 | 60.25 | 55.00 | 4.00 | 0.00 |
| Product length | 161 | | | | | | | | |

**B**

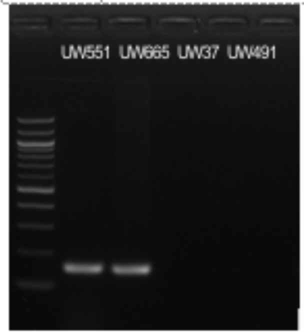


**C**

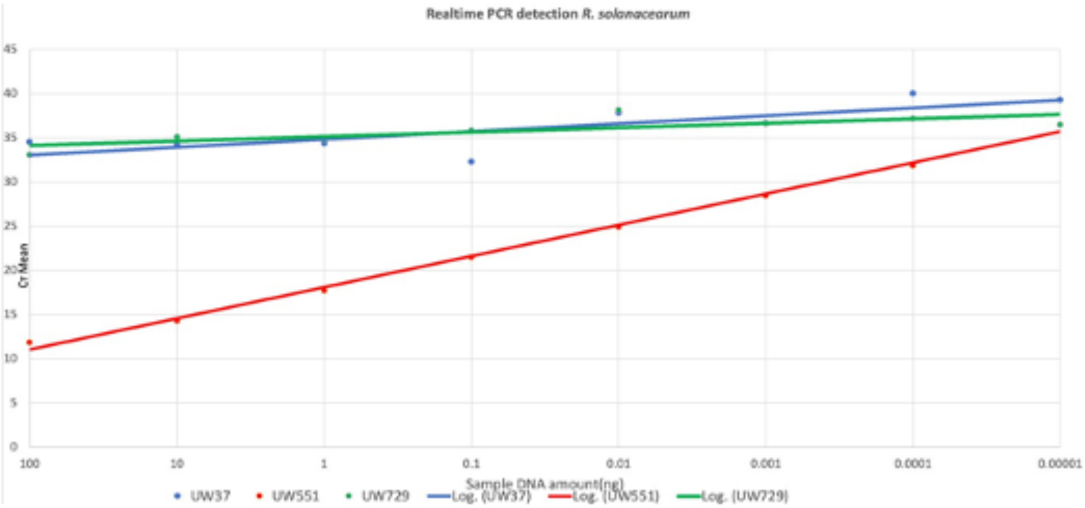


**Supplementary Figure S3**. Primer pair Seq1 was designed for conventional and quantitative PCR based on the sequence of gene RRSL_RS18155 of the *R. solanacearum* seq. 1 UW551 genome (Accession number GCA_002251655.1). **A** Primer sequences; **B** Gel image of products obtained by conventional PCR for sequevar 1 strains UW551 and UW 665 (both show a band of the correct size, as expected) and sequevar 2 strains UW 37 and UW491 (neither show any band, as expected); **C** Quantitative PCR result obtained for sequevar 1 strain UW551, sequevar 2 strain UW37, and sequevar 4 strain UW729 on an Applied Biosystems 7500 Realtime PCR System using a 20 µl reaction volume consisting of 10 µl PowerSYBR Green PCR Master Mix, 1 µl of primers (10µM), and genomic DNA: $10^2$ ng, $10^1$ ng, $10^0$ ng, $10^{-1}$ ng, $10^{-2}$ ng, $10^{-3}$ ng, $10^{-4}$ ng, $10^{-5}$ ng. Three technical replicates were performed; PCR cycle: Step1: 50°C, 2min, 95°C, 10 min, Step2: 40 cycles of 95°C, 15 sec + 60°C, 60 sec. Only UW551 (in red) gave a positive qPCR result, as expected.

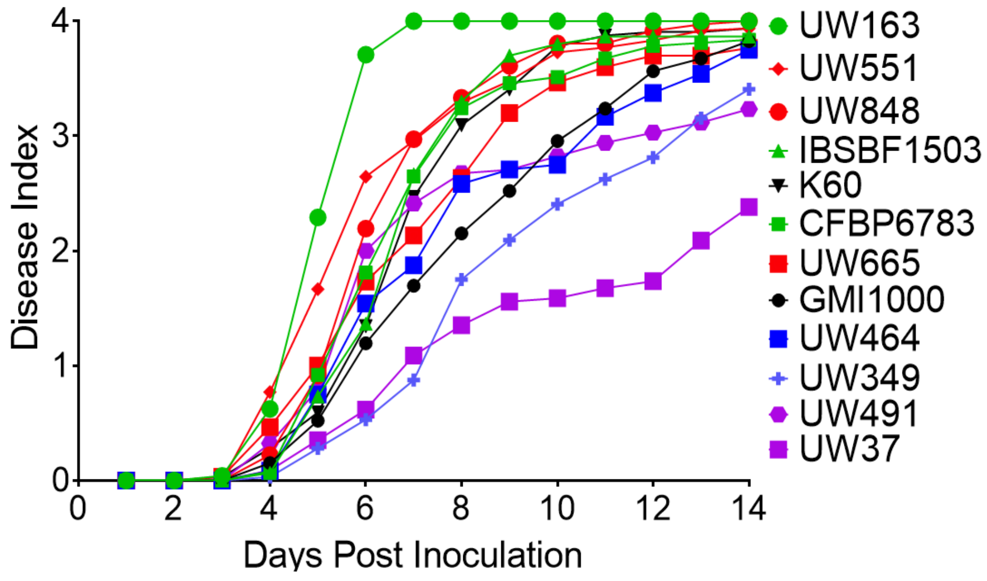


**Supplementary Figure S4. Virulence of selected RSSC strains on tomato at the generally permissive temperature of 28°C.** Disease progress curves over 14 days were generated for *Ralstonia* strains in seqs. 1, 2, 4, 28, and two non-phylotype IIB strains. Phylotype IIB strains are color-coded as follows: red, seq. 1; purple, seq. 2; blue, seq. 27 and 28; green, seq. 4; black, non-phylotype IIB strains. Each data point represents the mean disease index of 24-48 replicate plants in 4 to 8 independent experiments. These strains, having demonstrated the ability to kill more than half of tomato plants under experimental conditions at 28°C (corresponding to a final mean disease index >2.0), were designated as virulent and used for phenotyping analyses.

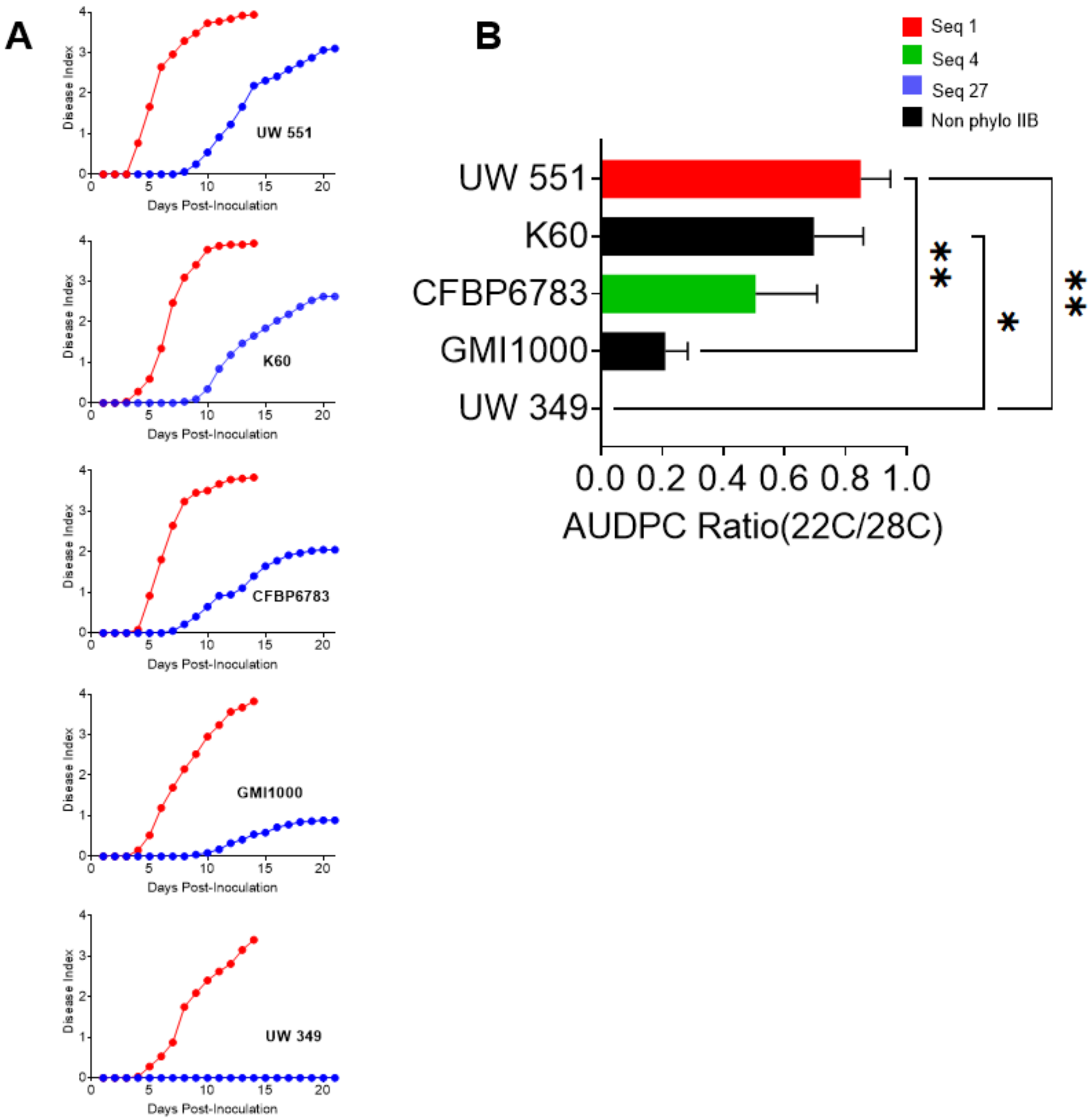


**Supplementary Figure S5. Virulence at 28°C on tomato does not predict virulence at 22°C on tomato for *Ralstonia* strains.** Unwounded 21-day-old tomato plants (cv. Bonny Best) were inoculated by pouring 50 ml of a $1x10^8$ CFU/ml water suspension of the indicated *Ralstonia* strain into the pot. Plants were incubated in a growth chamber at 28°C or 22°C and rated on a 0-4 disease index scale over 14 days at 28°C (red symbols) or 21 days at 22°C (blue symbols). **Disease progress curves of selected strains at 28°C (red symbols) and 22°C (blue symbols).** Each data point represents the mean disease index of 32 to 48 replicate plants in 5 to 8 independent experiments.

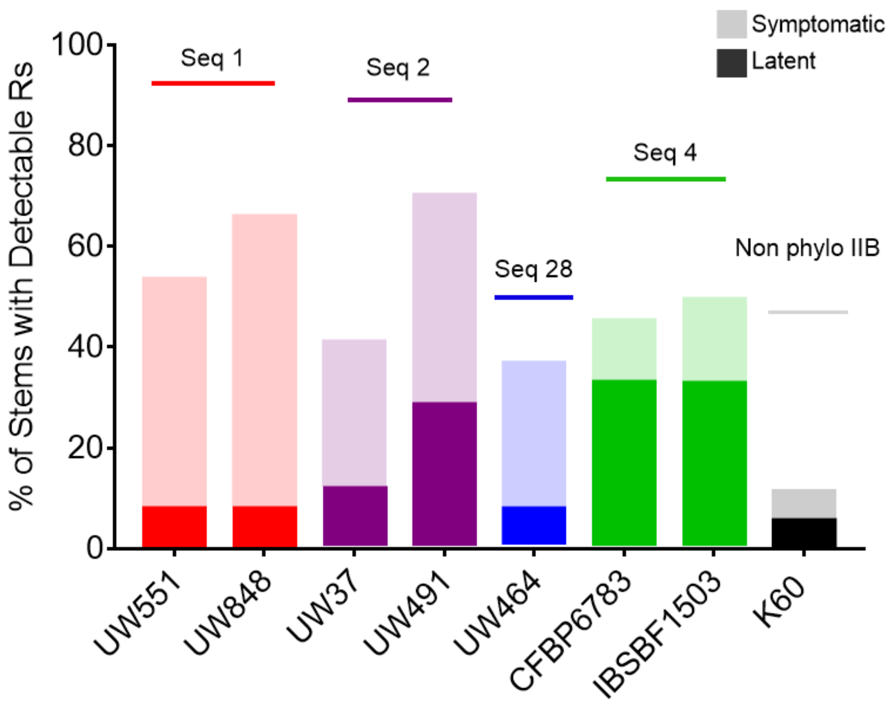


**Supplementary Figure S6: Frequency of detectable *R. solanacearum* cells in potato stems**. Each bar indicates the percentage of stems with detectable *R. solanacearum* populations from plants that were asymptomatic (darker lower bar) and symptomatic (lighter upper bar). Strains did not differ in their frequency of detection in potato stems ($P$=0.126, Pearson chi square test).
